# Disharmony of the world’s island floras

**DOI:** 10.1101/523464

**Authors:** Christian König, Patrick Weigelt, Amanda Taylor, Anke Stein, Wayne Dawson, Franz Essl, Jan Pergl, Petr Pyšek, Mark van Kleunen, Marten Winter, Cyrille Chatelain, Jan J. Wieringa, Pavel Krestov, Holger Kreft

## Abstract

**Aim:** Disharmony is a key concept in island biology that describes the imbalance in the representation of higher taxa on islands compared to their mainland source regions. Although there are strong theoretical arguments for the differential colonization success of different taxa on islands, the empirical evidence for disharmony remains largely anecdotal. Here, we develop a novel method for delineating island source regions and present the first global quantitative assessment of island disharmony.

**Location:** Global.

**Time period:** Recent.

**Major taxa studied:** Vascular plants.

**Methods:** We developed a generalizable method for estimating the source regions of an island flora based on statistical predictions of species turnover. We then designed two metrics to investigate disharmony from an island- and a taxon-centered perspective. First, we used linear mixed effects models to analyse the overall taxonomic bias of 305 island floras (compositional disharmony) as a function of geographical and climatic island features. Second, we applied linear models to examine the over- or under-representation of 450 vascular plant families on islands (representational disharmony) as a function of family size, age, higher taxonomic group and family-specific functional traits.

**Results:** We found that compositional disharmony scales positively with island isolation and negatively with island area, and is strongly modulated by climatic variables. In contrast, the relationship between representational disharmony and family-specific characteristics was weaker. We found significant effects of family species richness and pollination syndrome, whereas family age and all other tested functional traits remained without effect.

**Conclusions:** The taxonomic scope of the disharmony concept has historically limited its wider applicability, because higher taxa are inconsistent ecological proxies. However, our results provide a strong foundation for integrating disharmony with quantitative functional and phylogenetic approaches in order to gain a deeper understanding of assembly processes on islands.

## Introduction

Islands are renowned for their unique biotas, often characterized by high levels of endemism (Kier *et al.*, 2009), species radiations (Givnish *et al.*, 2009), relictual taxa (Cronk, 1997), or peculiar shifts in species’ life history and morphology (Carlquist, 1965). These features can be attributed to the isolated nature of islands (Weigelt & Kreft, 2013), which strongly affects the fundamental processes controlling species diversity: immigration, speciation, and extinction. Thus, research on island systems has stimulated many seminal contributions to evolutionary (Darwin, 1859; Wallace, 1881) and ecological theory (MacArthur & Wilson, 1963, 1967; Hubbell, 2001; Whittaker *et al.*, 2008). The island research by Carlquist (1965, 1967, 1974) is undoubtedly among these seminal contributions, providing substantial insights into processes such as long-distance dispersal or adaptive radiation, and inspiring island research to the present (Traveset *et al.*, 2015). In contrast to the strictly analytical approach of MacArthur & Wilson’s equilibrium theory of island biogeography (MacArthur & Wilson, 1967, 1963), Carlquist’s work offers mostly a natural history perspective on island assemblages. While this perspective does not allow for quantitative predictions of e.g. species richness, it represents a powerful framework for understanding qualitative features of island biotas such as taxonomic composition or morphological adaptations (Midway & Hodge, 2012). One such feature is the striking taxonomic “imbalance” of many island biotas– a phenomenon known as disharmony (Carlquist, 1974, 1965).

Island disharmony refers to the systematic over- or under-representation of higher taxa (e.g. families) in island biotas compared to their mainland source regions (Whittaker & Fernández-Palacios, 2007). It is the result of selective assembly mechanisms – dispersal filtering, environmental filtering and biotic filtering (Carlquist, 1974; Keddy, 1992; Weiher *et al.*, 2011; Kraft *et al.*, 2015) – acting with particular rigor in island systems, and thus permitting only a subset of the mainland flora to successfully colonize islands. The interplay between geographical setting and taxon-specific colonization success highlights two distinct aspects of island disharmony. On the one hand, the overall compositional imbalance of island floras relative to their mainland source regions (compositional disharmony) should reflect the impact of ecological filters during their assembly, and thus show predictable variation with island-specific characteristics such as isolation, area, climatic conditions, or geological origin. On the other hand, the selectivity of these filters should result in a predictable over- or under-representation of individual taxa on islands (representational disharmony) that is associated with taxon-specific attributes related to e.g. dispersal ability or environmental tolerances. Indeed, numerous studies demonstrate that community composition of island floras is strongly dependent on the geographical setting (Whittaker *et al.*, 2008; König *et al.*, 2017) and taxon-specific attributes (Burns, 2005; Olesen *et al.*, 2010).

While the theoretical underpinnings of island disharmony are well established, the concept itself has been applied inconsistently and currently lacks a quantitative basis. In particular, the specification of mainland source regions is not trivial and often very coarse (Bernardello *et al.*, 2006) and the taxonomic bias of island floras is usually illustrated by means of anecdotal evidence rather than objective quantitative measures (Francisco-Ortega *et al.*, 2010). In addition, there are only a few studies that have examined whether the over- or under-representation of taxa on islands is globally consistent (but see e.g. Kreft *et al.*, 2010 for ferns or Taylor *et al.*, 2019 for orchids), and whether representational deviations are linked to taxon-specific attributes that supposedly affect colonization success (but see e.g. Grossenbacher *et al.*, 2017 and Razanajatovo *et al.*, 2018). Consequently, the empirical evidence for island disharmony remains fragmentary.

Here, our main aim is to put the concept of island disharmony on a quantitative footing. First, we present a novel method for estimating island-specific source regions and develop two indices that quantify compositional and representational disharmony. Based on this, we examine disharmony from an island- and a taxon-centred perspective using a unique data set of 305 island and 611 mainland floras including a total of 225,053 species. To disentangle island- and taxon-specific drivers of disharmony, we analyse compositional disharmony as a function of island isolation, area, geological origin, and climatic conditions and representational disharmony as a function of species richness, age, higher taxonomic group and predominant functional characteristics of 450 plant families. In particular, we are interested in the importance of dispersal, environmental and biotic filtering in creating disharmonic island floras. If dispersal filtering is the dominant cause of disharmony (Carlquist, 1967, 1974), we would expect strong positive effects of isolation on compositional disharmony as well as a consistently positive effect of dispersal-related traits on representational disharmony. Alternatively, if environmental or biotic filtering processes play an important role (Carvajal-Endara *et al.*, 2017; Grossenbacher *et al.*, 2017), we expect to find a strong response of compositional and representational disharmony to island climatic variables and pollination or competition-related traits, respectively.

## Methods

We examined the phenomenon of island disharmony from both an island- and a taxon-centred perspective (Figure 1). First, we assessed compositional disharmony, i.e. the phenomenon of island floras being taxonomically biased compared to their mainland source regions. Second, we investigated representational disharmony, i.e. the role of individual taxa in creating compositional disharmony by assessing their global representation on islands compared to the mainland. In both cases, we chose families as the focal taxonomic level. Given that disharmony explicitly deals with the representation of higher taxonomic groups on islands, families provide a reasonable compromise between ecological uniformity and taxonomic aggregation. In contrast, higher taxonomic levels such as orders encompass too many species that are too heterogeneous in their attributes to be ecologically meaningful study units, whereas lower levels such as genera are too numerous to be harmonically represented in any island flora.

**Figure 1:**
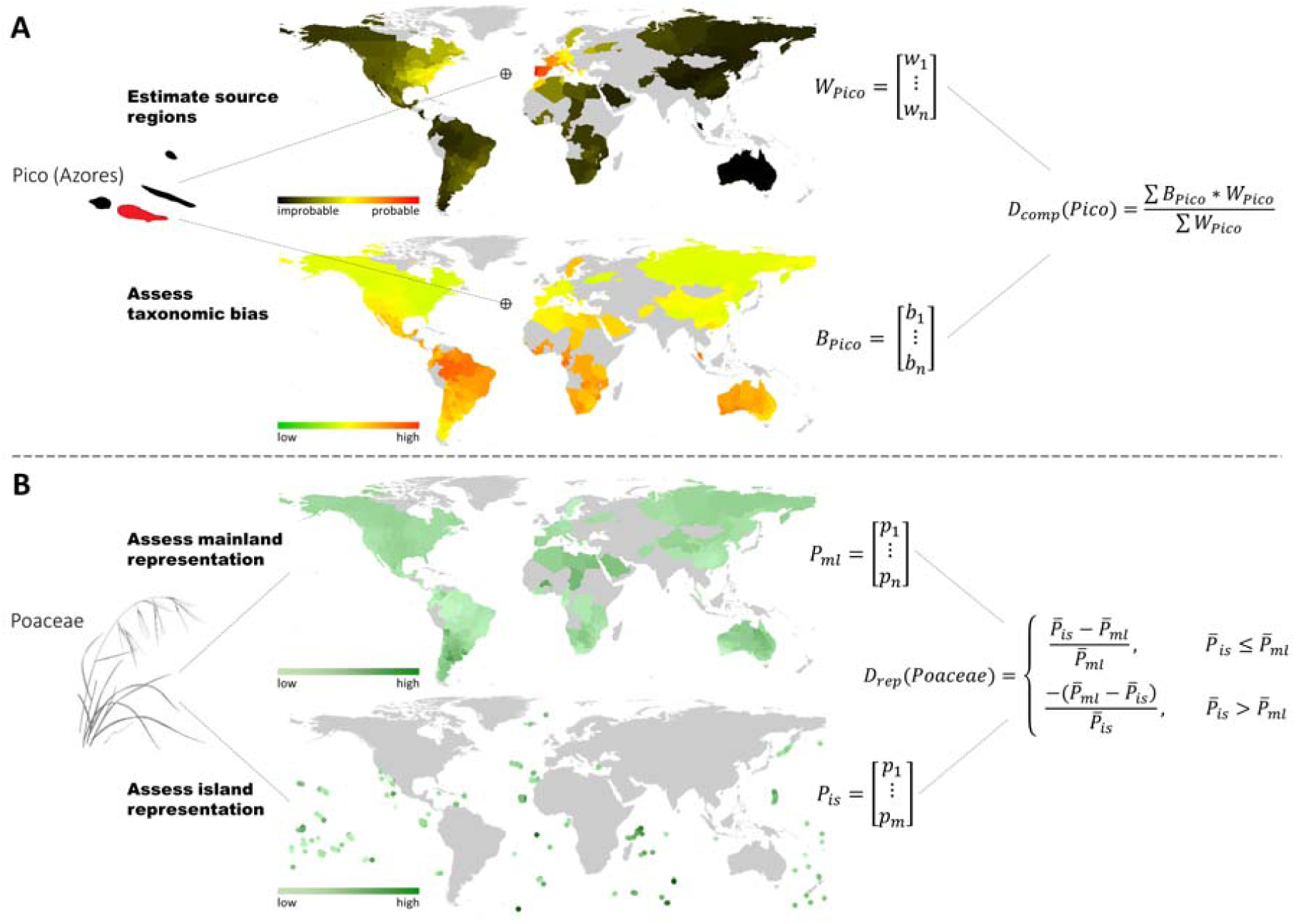
Schematic representation of the quantification of compositional and representational disharmony. (A) Calculation of compositional disharmony by the example of Pico Island (Azores). Source regions were estimated based on predictions of species turnover between the focal island and all mainland units (W_Pico_, upper plot). The taxonomic bias between the focal island and all mainland units was quantified using Bray-Curtis dissimilarity on relative proportions of plant families (B_Pico_, lower plot). The compositional disharmony of the focal island (D_comp_) was then calculated as the mean taxonomic bias relative to all mainland regions, weighted by their respective source region weight. (B) Estimation of representational disharmony by the example of Poaceae. Representational disharmony was quantified based on the mean proportion of the focal taxon in mainland floras 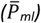 and island floras 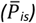. The corresponding index (D_rep_) transforms the ratio between these two components to a range between −1 (family occurs on the mainland only) and 1 (family occurs on islands only).

### Data collection

All ecological and environmental data were obtained from the Global Inventory of Floras and Traits database (GIFT, Weigelt *et al.*, 2019), a comprehensive resource for macroecological analyses of global plant diversity. The primary data type in GIFT are regional plant checklists that are integrated with additional information at the level of taxa (e.g. functional traits, taxonomic placement, phylogenetic relationships, floristic status) and geographical units (e.g. climatic and environmental conditions, socioeconomic factors, physical geographic properties). A detailed description of the database structure and data processing workflows related to taxonomic standardization, functional traits and geographic information is available in Weigelt *et al.* (2019).

For this study, we used only species checklists that report the floristic status of all listed species and excluded non-native species occurrences from the analyses. Checklist completeness was evaluated based on the reference type (e.g. multi-volume Floras being more reliable than rapid assessments), the general text included in the reference (e.g. statements regarding sampling effort, timeframe, or use of additional data sources), and general properties of the species list (e.g. plausible number of species for the given area and biome, species-to-genus ratio, presence of regionally important taxa). Checklists with considerable deficits in either of these categories were excluded. We then combined checklists referring to the same geographical unit and removed all geographical units that were not covered by either a checklist of vascular plants or by several checklists that cover all divisions of vascular plants (e.g. separate lists for pteridophytes and seed plants). The resulting dataset contained native vascular plant checklists for 611 mainland and 305 islands units (see data references). For each geographical unit, we calculated climatic characteristics based on median values of CHELSA climate layers (Karger *et al.*, 2017). We chose mean annual temperature (Bio1), mean annual precipitation (Bio12), temperature seasonality (Bio4) and precipitation seasonality (Bio15) as widely established measures of energy and water availability. For islands, we additionally calculated maximum elevation as a measure of habitat heterogeneity and researched the geological origin and corresponding archipelago (where applicable) in pertinent literature.

We obtained the total number of species per family from The Plant List (The Plant List, 2013) and supplemented it with values from Christenhusz & Byng, 2016) where The Plant List did not resolve the respective family. The assignment of taxonomic supergroups (distinguishing between angiosperms, gymnosperms and pteridophytes) and family ages was based on a recently constructed megaphylogeny of vascular plants (Qian & Jin, 2016). For seed plants only, we selected five functional traits that reflect dispersal ability (seed mass, dispersal syndrome [anemochorous, autochorous, hydrochorous, zoochorous, unspecialized]), life history (woodiness [woody, non-woody], plant height), and reproductive characteristics (pollination syndrome [abiotic, biotic]) of a family. Trait data were obtained through literature research and the aggregation of species-level information available in GIFT. For categorical traits, we assigned a value if multiple taxonomic literature resources (Kubitzki, 1990-2014; Vamosi & Vamosi, 2010; Hawkins *et al.*, 2011; Hintze *et al.*, 2013) indicated the same predominant trait syndrome for a family (woodiness: 289 families, dispersal syndrome: 117 families, pollination syndrome: 229 families). If information from taxonomic resources was missing or conflicting for a particular family–trait combination, we assigned a value if > 66 % of the species-level information in GIFT was identical, i.e. a predominant trait syndrome within the family could be inferred from species-level data (woodiness: 174 families from 100,028 species, dispersal syndrome: 156 families from 3,667 species, pollination syndrome: 103 families from 2,993 species). Otherwise, we assigned no value. For numerical traits, we calculated the median value per family based on available species-level data in GIFT (seed mass: 340 families from 23,867 species, plant height: 384 families from 53,449 species).

### Compositional disharmony

#### Quantification

The quantification of compositional disharmony requires estimates of island-specific source regions as well as an objective measure of taxonomic bias (Figure 1A). We based our method for estimating source regions on the fact that geographic distance and environmental gradients produce distinctive and predictable patterns in species turnover (Fitzpatrick *et al.*, 2013; König *et al.*, 2017). Species turnover is a richness-insensitive measure of compositional similarity that quantifies the proportion of shared species between assemblages (Baselga, 2010). This makes turnover a crucial concept for constructing biogeographical species pools and delineating source regions (Carstensen *et al.*, 2013).

In contrast to existing approaches for the reconstruction of biogeographical source regions (e.g. Graves & Rahbek, 2005; Papadopulos *et al.*, 2011), our method is based on statistical predictions (rather than pairwise calculations) of species turnover. Consequently, our framework does not require floristic data for the focal area or any of the potential source regions, but only a fitted model of species turnover and a set of environmental predictor variables. We used generalized dissimilarity modelling (GDM, Ferrier *et al.*, 2007) to model species turnover (β_sim_, Koleff *et al.*, 2003) as a function of geographic distance and differences in mean annual temperature, mean annual precipitation, temperature seasonality and precipitation seasonality. These covariates have emerged as strong predictors of large-scale species turnover of vascular plants (König *et al.*, 2017). We fitted the model using species checklists of mainland units only (n = 611, deviance explained = 80.5%), because island floras exhibit strong imprints of ecological filtering, which would mask the very effects we aim to quantify in this study. We then used the fitted model to predict species turnover between each island (n = 305) and a global set of 6495 equal-area grid cells based on geographical and environmental information only. To relate these island-specific predictions of species turnover back to the floristic data available in GIFT, we aggregated the results by calculating the area-weighted mean of grid values per mainland unit (Figure 1A, upper plot). The resulting *m* × *n* matrix, **W** (for weight), contained the statistically expected proportion of shared species (1-β_sim_) between all *m* islands and all *n* mainland units from a “mainland perspective”, i.e. assuming that island floras assemble under the same conditions as floras on the mainland. The matrix W can thus be interpreted as reflecting the importance of a given mainland unit as a source region for a given island, while excluding the effects of modified filtering during island colonization. We set W_m,n_ = 0 for mainland units with very low values, i.e. highly improbable source regions for a given island, to ensure a balanced estimation of compositional disharmony (see SI Text 1 for details). To validate our method, we compared the results with empirical source region reconstructions based on floristic or phylogenetic relationships between island and mainland floras.

To assess the actual imbalance in the representation of higher taxa on islands relative to the mainland, we calculated the relative proportion of plant families for all geographical units. Based on these proportions, we computed the Bray-Curtis dissimilarity for all pairwise island-mainland combinations (Figure 1A, lower plot), yielding an *m × n* matrix, **B** (for bias). The Bray-Curtis index is the abundance-based version of the Sørensen index that relates the summed differences between two variables to their overall sum. Although usually applied to count data, the index also works with relative proportions (Greenacre & Primicerio, 2013), making it an appropriate measure of taxonomic bias.

Finally, we calculated the compositional disharmony of each island, **D**_comp_, as the mean taxonomic bias of a given island i relative to all mainland units (B_i_), weighted by their respective importance as a source region for the specific island (W_i_). D_comp_ ranges between 0 and 1, with higher values indicating more disharmonic floras (Figure 1A).

#### Analysis

We log_10_-transformed island area, distance to the mainland, mean annual precipitation, temperature seasonality, and precipitation seasonality because of strongly skewed distributions. We then scaled all continuous predictor variables to µ = 0 and σ = 1 to obtain standardized model coefficients. The geological origin of islands was considered as a categorical variable (distinguishing between volcanic islands, atolls, islands formed by tectonic uplift, continental fragments, and shelf islands; adopted from König *et al.*, 2017).

To quantify the effects of geographical and climatic variables on compositional disharmony (D_comp_), we fitted generalized mixed effects models within a Bayesian framework (brms R-package, Bürkner, 2017). We used models from the beta family with a log-link function because our index for compositional disharmony is a proportional value ranging between 0 and 1. Following Bunnefeld & Phillimore (2012), we specified archipelago as random effect and allowed for random intercepts, but not for random slopes. In addition, we fitted a model without random effects to evaluate the unique explanatory power of the fixed effects representing climate and geography.

### Representational disharmony

#### Quantification

In any flora, a given family accounts for a proportion of species ranging between 0 and 1. To quantify representational disharmony, we calculated the mean proportion of each family across all mainland units 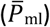 and oceanic islands (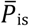, see Figure 1B). For the calculation of 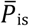, we focused on oceanic islands (volcanic islands, islands formed by tectonic uplift, atolls), because their floras reflect the effects of dispersal, environmental and biotic filtering most clearly (Whittaker & Fernández-Palacios, 2007) and thus allow for the most rigorous assessment of taxon-specific drivers of disharmony.

To obtain an index of representational disharmony (D_rep_) for a given family, we then transformed the ratio between 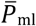 and 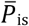 according to the formula given in Figure 1B. The index is symmetric and obtains values ranging from −1 to 1, with the sign indicating whether the focal family is proportionally more common in mainland floras (negative values) or in island floras (positive values; SI Figure 1). For example, a family with D_rep_ = 0.5 has, on average, a two times higher proportion in island floras than in mainland floras, whereas a family with D_rep_ = −0.9 has a 10-times higher proportion on the mainland. The special cases of D_rep_ = −1, D_rep_ = 0 and D_rep_ = 1 indicate a family’s restriction to the mainland, equal proportional representation on islands and the mainland, and restriction to islands, respectively.

#### Analysis

To analyse the effects of family characteristics on representational disharmony (D_rep_), we used linear regression models fitted within a Bayesian framework (brms R-package, Bürkner, 2017). We fitted a multi-predictor model with family species richness, family age and taxonomic supergroup (distinguishing between angiosperms, gymnosperms and pteridophytes) as predictors, but had to omit all functional trait variables because missing data points would have drastically reduced the sample size. Instead, we analysed the impact of functional traits on representational disharmony (D_rep_) with single-predictor linear models.

To test for phylogenetic signal in D_rep_, i.e. whether closely related taxa tend to be similarly over- or under-represented on islands, we calculated Abouheif’s C_mean_ using the *phylosignal* R-package (Keck, 2015). The C_mean_ index is a measure of phylogenetic autocorrelation that quantifies the squared differences between values (in this case D_rep_) of neighbouring tips in a phylogeny (Münkemüller *et al.*, 2012). The index ranges from −1 to 1, with values further away from 0 indicating negative and positive autocorrelation, respectively.

All analyses were performed in the R statistical programming language, version 3.5.2 (R Core Team, 2019).

## Results

### Source region estimation

Our method for statistically estimating island source regions showed strong agreement with empirical source region reconstructions (Figure 2). Accordingly, most island floras are derived from a limited set of nearby and climatically similar mainland regions. The estimated source regions for the Falkland Islands, for instance, are highly concentrated in the nearby non-tropical parts of South America, which corresponds closely to the account given by Moore (1968) (Figure 2B). Similarly, the most likely source regions for the flora of Lord Howe or Cocos Island are restricted to a few regions in Australasia and the Neotropics, respectively (Figure 2D-E). With increasing isolation from the mainland, however, the distribution of island source regions becomes more diffuse in both the statistical and empirical reconstructions (Figure 2A, C, F). For example, a wide, circum-Pacific distribution of source regions for the Hawaiian flora emerges in our statistical estimates as well as phylogenetic analyses of Price & Wagner (2018) (Figure 2A). While the accuracy of our estimates seems to decrease slightly for very isolated islands (e.g. Figure 2F), the overall congruency of our statistically estimated source regions with empirical studies demonstrates the robustness of our method.

**Figure 2:**
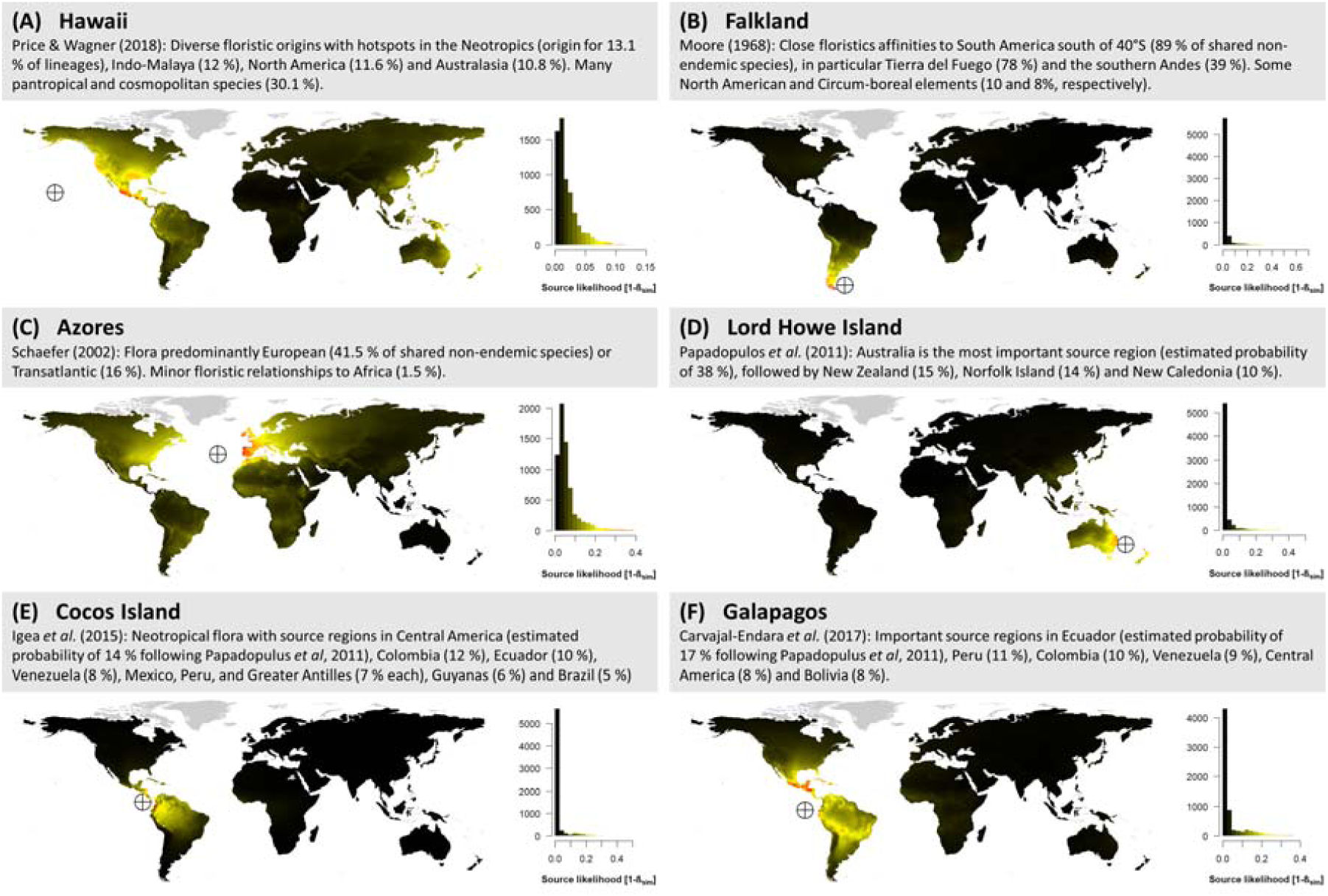
Exemplary comparison of empirically reconstructed and statistically modelled source regions for six islands. Grey boxes summarize the results of previous source region reconstructions based on floristic or phylogenetic affinities. Maps show corresponding statistical source region estimates (proportion of shared species, 1-β_sim_) between the focal island and 6495 equal-area grid cells (∼23,300 km^2^ each). Predictions were derived from a generalized dissimilarity model (Ferrier et al., 2007) fitted with geographical and climatic characteristics of 611 mainland floras worldwide. Histograms show the distribution of predicted values for each focal island (note that the range of values differs among islands).

### Compositional and representational disharmony

Compositional disharmony (D_comp_) ranged between 0.19 (Corsica, Mediterranean Sea) and 0.87 (Clipperton Island, East Pacific). Overall, the most harmonic island floras were found in the Mediterranean Basin and off the shores of temperate continental regions (East Asia, Europe, North America). Particularly disharmonic floras were located at high latitudes and in the Central Pacific (Figure 3).

**Figure 3:**
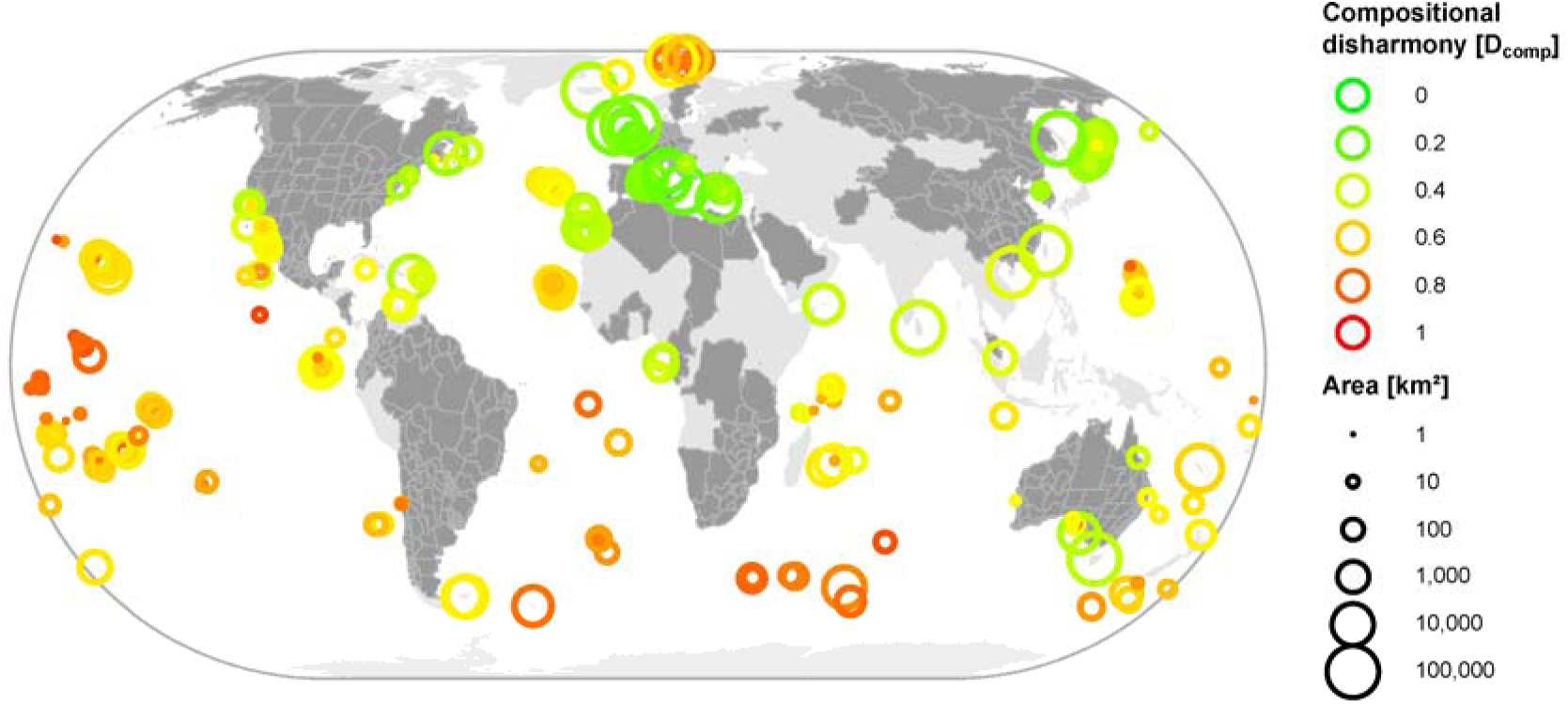
Compositional disharmony of 305 island floras worldwide. Mainland regions that were used for source pool estimation and floristic comparisons (n = 611) are coloured in dark grey.

In agreement with our expectations, compositional disharmony increased with island isolation and decreased with island area (Figure 4). Maximum elevation had a negative effect on compositional disharmony, i.e. topographically flat islands tended to harbour more imbalanced floras. Also climatic variables had consistently negative and unexpectedly strong effects. Measured by the slope of standardized regression coefficients temperature seasonality was the most important predictor of compositional disharmony, and the effect size of mean annual temperature was comparable to that of island area (islands with cold, seasonal climates having more disharmonic floras, Figure 4). After controlling for the effects of island area, elevation, isolation, and climate in the multi-predictor setting, the geological origin of islands did not show strong effects – only atolls featured significantly more disharmonic floras than other island types. The overall model fit was very high with a Bayesian R^2^ of 0.90 (full model including random effects) and of 0.58 (fixed effects only).

**Figure 4:**
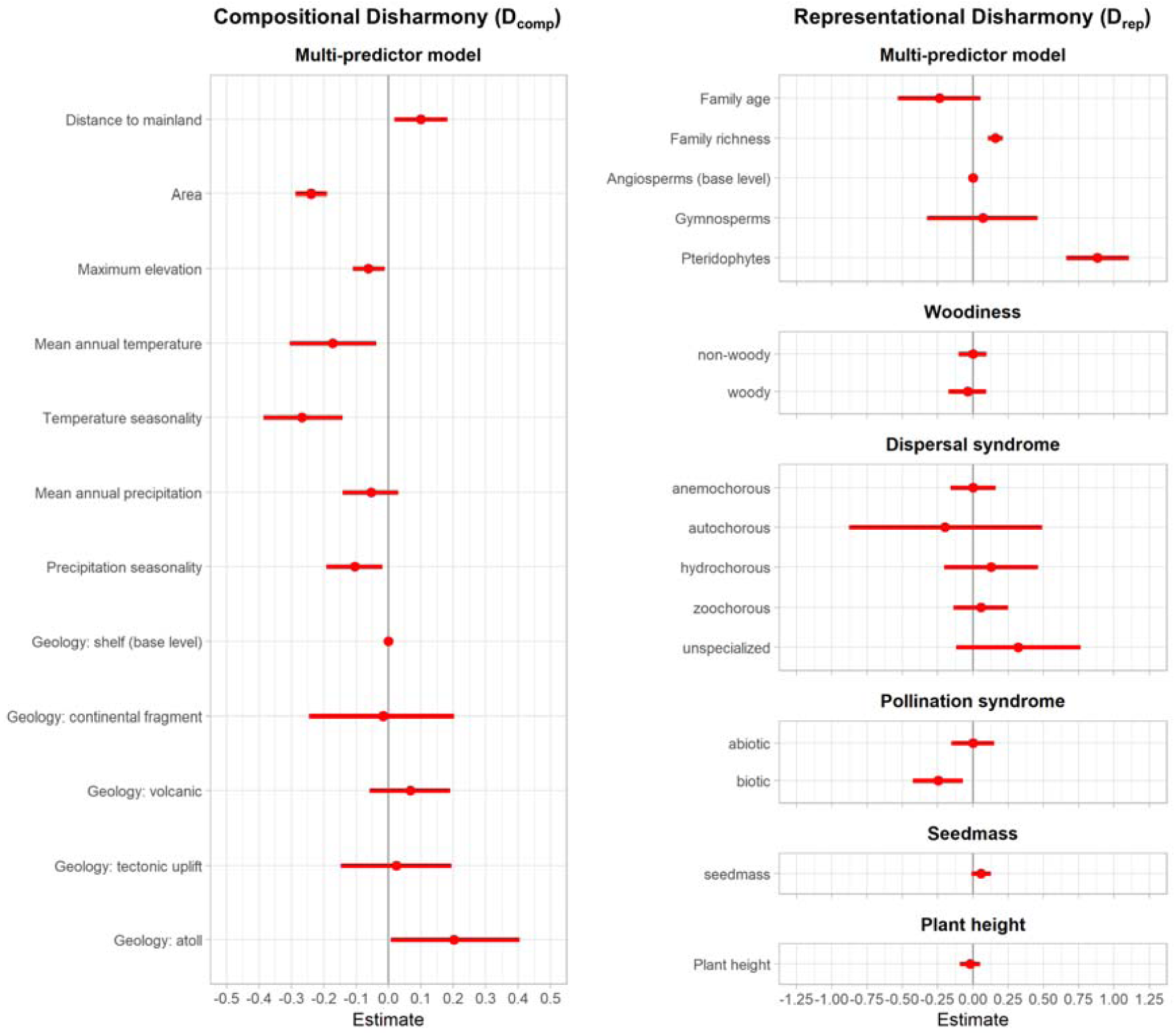
Standardized parameter estimates (dot) and 95% Bayesian credible intervals (whiskers) for potential drivers of compositional disharmony (left) and representational disharmony (right). Estimates for island geological origin and taxonomic group are relative to shelf islands and Angiosperms, respectively.

Representational disharmony varied widely among plant families. Most notably, pteridophyte families were greatly over-represented on islands, whereas angiosperm and especially gymnosperm families were mostly under-represented (see SI Data). A few families such as Cycadaceae (gymnosperms, D_rep_ = 0.62) or Marsileaceae (pteridophytes, D_rep_ = −0.70) deviated from this general pattern. The multi-predictor model confirmed the over-representation of pteridophyte families on islands relative to angiosperm and gymnosperm families (Figure 4). Species richness was positively associated with the representation of families on islands, whereas family age had marginal negative effects. However, the explanatory power of the model was relatively low (Bayesian R^2^ = 0.19). The single predictor models of family-level functional traits did not reveal strong effects either. We did not find significant effects of life history traits (growth form, plant height) or dispersal traits (dispersal syndrome, seed mass). Only for pollination traits did we significant effect, namely a slight over-representation of families with abiotic pollination syndromes over those with biotic pollination syndromes (Figure 4).

In agreement with the generally weak predictive power of family level characteristics for representational disharmony, we did not find a significant phylogenetic signal in D_rep_ for seed plants (C_mean_ = 0.05, p = 0.065), but only for all vascular plants combined including the strongly over-represented pteridophyte clade (C_mean_ = 0.16, p = 0.001, SI Figure 3).

## Discussion

Our results show that compositional disharmony, i.e. the imbalance of an island flora relative to its mainland source regions, is a common feature of islands worldwide. The degree of compositional disharmony is well explained by the classical biogeographical variables of isolation and area in conjunction with additional island characteristics such as elevation, geological origin and climate. We found less clear relationships between family-specific characteristics and representational disharmony, i.e. the proportional over- or under-representation of individual families on islands. The most important predictor of representational disharmony was a simple categorization of families into angiosperms, gymnosperms and pteridophytes. Species richness and pollination traits showed weak but significant effects, but we found no evidence for effects of dispersal and life history traits.

One innovation of the present study is the development of a method for estimating biogeographical source regions based on statistical predictions of species turnover. Existing methods typically infer source regions by comparing the composition of species, genera or families (e.g. Papadopulos *et al.*, 2011; Moore, 1968) or the phylogenetic relationships in the focal region with that of several potential source regions. The broad geographical scope of the floristic literature underlying such comparative analyses only allows for the delineation of relatively coarse source regions such as continents, biogeographical regions or countries. However, a more fine-grained understanding of potential source regions is often needed. More recent methods therefore derive the compositional structure of any given location by means of so-called assemblage-dispersion fields, which are stacked geographical ranges of all species occurring in the focal region (Carstensen *et al.*, 2013; Graves & Rahbek, 2005). Although assemblage dispersion fields are theoretically capable of providing high-resolution estimates of biogeographical source regions, they practically require knowledge on the complete distribution of all examined species. This is still beyond reach for many regions and taxa (Hortal *et al.*, 2015). Moreover, like all methods that are based on contemporary species distributions or assemblages, they break down when the focal region features high rates of endemism, which is particularly important for island systems (Kier *et al.*, 2009). The statistical approach we outlined in this study avoids the above-mentioned limitations because it does not depend on floristic data for any of the investigated geographical regions. Instead, it depends on how well the statistical model and the input data (which may come from other floras) capture the general drivers underlying large-scale species turnover. This makes it considerably less data-intensive than alternative species-level approaches (e.g. Graves & Rahbek, 2005), yet offers much finer spatial resolutions than checklist-based methods (e.g. Papadopulos *et al.*, 2011).

The assembly of an island flora is an accumulative process, acting over millions of years. The extended temporal dimension introduces uncertainties to the estimation of island source regions because the climate, habitat distribution, position, size and shape of both islands (Whittaker *et al.*, 2008; Weigelt *et al.*, 2016) and source regions (Galley & Linder, 2006; Pokorny *et al.*, 2015) may have changed considerably since colonization. Cronk (1987) illustrates this for the flora of Saint Helena, which is mostly derived from a now-extinct wet forest flora that occupied large parts of Southern Africa during the Miocene. Moreover, the effective isolation of an island is difficult to quantify and depends not only on the distance to the mainland, but also on the availability of stepping stones and the direction of predominant sea and wind currents (Cook & Crisp, 2005; Weigelt & Kreft, 2013), as well as the dispersal abilities of the focal taxon. Even different habitats or elevational zones within an island may sample from distinct source regions on the mainland, and thus vary in their degree of isolation (Steinbauer *et al.*, 2012). Thus, many aspects affect the accuracy of biogeographical source region reconstructions in general, but may be particularly relevant for statistical approaches that do not involve direct comparisons of species assemblages. However, our validation revealed high levels of congruency between empirical and statistical source region reconstructions (Figure 2), and further methodological refinements and validations may help to find the appropriate balance between complexity and utility. Considering that quantitative estimates of source regions are still lacking for most island floras, statistical reconstructions might add a promising new tool to island biogeographical research.

Dispersal filtering has long been regarded as the predominant process in the assembly of island biotas, and therefore the main driver of disharmony (Carlquist, 1966, 1967). Our results reveal that this is only partly true. On the one hand, the strong effect of isolation on island disharmony indeed suggests a major role of dispersal filtering in removing less dispersive taxa from the set of potential colonizers of an island. All gymnosperms except for Araucariaceae and Cycadaceae were under-represented on oceanic islands, which seems to fit the classical notion of gymnosperms as poor dispersers. Likewise, pteridophytes – possessing superior long-distance dispersal capabilities via ultra-light spores – were found to be strongly over-represented in island floras. These findings are in line with previous studies, which interpreted these broad taxonomic patterns as the outcome of selective dispersal filtering (Kreft *et al.*, 2010; Rumeu *et al.*, 2014; Weigelt *et al.*, 2015). On the other hand, the strong effects of area and climatic variables, as well as the relationship between representational disharmony and pollination- but not dispersal-related traits, indicate an important role of non-dispersal related processes. Pollination is increasingly recognized as a critical factor for the colonization of islands (Olesen *et al.*, 2010; Alsos *et al.*, 2015; Grossenbacher *et al.*, 2017). Given the general scarcity of animal pollinators on islands, abiotic pollination syndromes and the ability to self-pollinate should provide an advantage over biotic pollination (especially when pollinator-specific) or strict outcrossing (Baker, 1955; Razanajatovo *et al.*, 2018). Indeed, we found families with predominantly abiotic pollination syndromes (wind pollination, water pollination) to be proportionally slightly more common on islands than families that rely on biotic pollination (insects, bats, birds, etc.)(Figure 4). However, all other family-level traits had no detectable effects on representational disharmony. The lacking effect of classical dispersal traits such as seed mass, dispersal syndrome or fruit type seems to contradict longstanding assumptions regarding their relevance for island colonization (Carlquist, 1974; Howe & Smallwood, 1982; Portnoy & Willson, 1993). Indeed, while a relationship between such traits and dispersal distance is evident at small scales up to a few kilometers (Tackenberg *et al.*, 2003), long-distance dispersal seems to operate with such high levels of stochasticity that this relationship vanishes (Higgins *et al.*, 2003; Nathan, 2006; Nogales *et al.*, 2012). Moreover, many large-seeded species that are adapted to dispersal by birds or seawater are successful long-distance dispersers, defying the common notion of small seeds as indicator of good dispersibility. Other studies even find no relationship at all between dispersal traits and colonization success (Heleno & Vargas, 2014).

Abiotic factors such as climate or soil substrate also act as filters for colonizing plant species, as recently demonstrated for the Galapagos Islands (Carvajal-Endara *et al.*, 2017) and New Caledonia (Isnard *et al.*, 2016). The strong effect of temperature-related variables in our analyses seems to support (abiotic) environmental filtering as an important driver of island disharmony. However, the climatic variables in our models did not reflect climatic similarity to the mainland (which would be a plausible cause of disharmony, but was accounted for during source region estimation), but the average conditions of the islands themselves. Potential explanations for the negative relationship between compositional disharmony and island temperature and temperature seasonality include (1) stronger environmental filtering on islands with temperate or polar climates, (2) higher speciation rates on warm tropical islands, or (3) glacial dynamics limiting the available time for colonization on cold, high-latitude islands. In addition, separating abiotic and biotic drivers of community assembly is often difficult because competitors may preclude the establishment of colonizing species in generally suitable habitats, thus creating artificial environmental gradients in the composition of communities (Kraft *et al.*, 2015).

Even though the above considerations provide plausible explanations for the role of ecological factors in creating island disharmony, we want to stress that compositional disharmony is affected by neutral sampling effects related to species richness. Island floras can never be a perfect compositional representation of the much larger pool of mainland species, because the number of families on an island is constrained by the number of species. Thus, extremely small proportions that arise, for example, in the case of rare families in species-rich mainland floras cannot be reproduced on islands, which inevitably increases compositional disharmony with decreasing species number (SI Figure 4). This interpretation moreover implies that compositional disharmony is also dependent on the size and spatial extent of the mainland source pool, as larger source pools usually include a higher number of taxa and are thus less likely to be represented harmonically in an island flora. This dependency might provide a further piece in the puzzle of understanding the disharmonic floras of highly isolated islands such as Hawaii or the Azores (see Figure 2B,C), whose source regions often encompass different biogeographical regions and continents.

If the overall bias of island floras is rather accurately predicted by geographical and climatic island features, why does the representation of individual families on islands seem so unrelated to their functional traits? We consider two aspects to be relevant here. First, if the taxonomic literature did not indicate a predominant trait syndrome within a family, we attempted to approximate it based on species-level information of varying completeness (see Methods section and SI Figure 2). Missing data is a common problem in trait-based ecology (Taugourdeau *et al.*, 2014; Penone *et al.*, 2014) and a major source of uncertainty and bias in the characterization of ecological patterns (Hortal *et al.*, 2015). We therefore caution that our findings on representational disharmony depict trends rather than a definitive assessment. Second, a taxonomic perspective – especially when focusing on a fixed taxonomic level – is not an ideal framework for examining the outcomes of the complex ecological processes that produce disharmonic island floras. In the scientific literature, examples of disharmonic floristic elements range from small genera (e.g. *Metrosideros* in Carlquist, 1966) to large taxonomic clades (e.g. pteridophytes in Braithwaite, 1975). Some studies assemble several such examples for a particular island or island group in order to arrive at a more general conclusion (Carlquist, 1967; Whittaker *et al.*, 1997). In fact, this is a reasonable approach because the degree to which taxa are consistently over- or under-represented on islands depends on their uniformity in terms of colonization success, and thus in terms of dispersal abilities, environmental tolerances and degree of biotic specialization. These parameters may vary greatly even within small families (Howe & Smallwood, 1982), but on the other hand show remarkable consistency within large taxonomic groups (Farjon, 2010), such that the required taxonomic level of observation is variable. Consequently, the classical concept of island disharmony *sensu* Carlquist (1965, 1974) generalizes poorly across taxonomic groups.

We demonstrated that disharmony is a widespread feature of island floras worldwide, and that the traditional concept can be put on a quantitative footing. The generality and predictive power of the concept of island disharmony has historically been limited by its focus on taxonomic groups. However, given the rapid advances in ecological data availability and analytical tools, the approximation of ecological characteristics by means of taxonomic affiliation will eventually become obsolete. Instead, the assembly mechanisms that disharmony aims to reflect and explain can be investigated more directly using quantitative methods that are informed by functional and phylogenetic data. The outlined approach for the estimation of island source regions as well as the proposed metrics for quantifying island disharmony provide a big step towards this objective, and offer robust empirical insights into the factors shaping the composition of island floras.

## Acknowledgments

JP and PP were supported by EXPRO grant no. 19-28807X (Czech Science Foundation) and long-term research development project RVO 67985939 (The Czech Academy of Sciences).

## Data availability

All relevant data are available as supporting data items.

## Data references

|  | checklists | traits |
| --- | --- | --- |
| 3D Environmental (2013). *Profile for management of the habitats and related ecological and cultural resource values of Boigu Island* | 0 | 1 |
| Acevedo-Rodríguez, P. & Strong, M.T. (2007). Catalogue of the seed plants of the West Indies Website, Available at: http://botany.si.edu/antilles/WestIndies/catalog.htm (last accessed: March 01, 2011) | 1 | 0 |
| Alves, R.J.V. (1998). *Ilha da Trindade & Arquipélago Martin Vaz. Um ensaio geobotânico*. Serviço de Documentação da Marinha, Rio de Janeiro, Brasil | 1 | 0 |
| Arakaki, M. & Cano, A. (2003). Composición florística de la cuenca del río Ilo-Moquegua y Lomas de Ilo, Moquegua, Peru. *Revista Peruana de Biología*, **10**, 5–19 | 0 | 1 |
| Arechavaleta, M., Rodríguez, S., Zurita, N. & García, A. (2009). *Lista de especies silvestres de Canarias. Hongos, plantas y animales terrestres*. Consejería de Medio Ambiente y Ordenación Territorial, Gobierno de Canarias, Santa Cruz de Tenerife, Spain | 1 | 0 |
| Ashmole, P. & Ashmole, M. (2000). *St Helena and Ascension Island. A natural history*. Anthony Nelson Ltd, Oswestry, Shropshire, UK | 1 | 0 |
| Athens, J.S., Blinn, D.W. & Ward, J.V. (2007). Vegetation history of Laysan Island, Northwestern Hawaiian Islands. *Pacific Science*, **61**, 17–37 | 1 | 0 |
| Baker, M.L. & Duretto, M.F. (2011). *A census of the vascular plants of Tasmania*. Tasmanian Herbarium, Tasmanian Museum and Art Gallery, Hobart, Australia | 1 | 0 |
| Barker, W.R., Barker, R.M., Jessop, J.P. & Vonow, H.P. (2005). Census of South Australian vascular plants. *Journal of the Adelaide Botanic Gardens Supplement*, **1**, 1–396 | 1 | 0 |
| Belhacene, L. (2010). Catalogue 2010 des plantes vasculaires du département de la Haute-Garonne. *Supplément à Isaatis*, **10**, 1–145 | 1 | 0 |
| Benito, B.M., Lorite, J., Pérez-Pérez, R., Gómez-Aparicio, L., Peñas, J. & Robertson, M. (2014). Forecasting plant range collapse in a mediterranean hotspot. When dispersal uncertainties matter. *Diversity and Distributions*, **20**, 72–83 | 0 | 1 |
| Bernal, R., Gradstein, S.R. & Celis, M. (2015). Catálogo de plantas y líquenes de Colombia, Available at: http://catalogoplantasdecolombia.unal.edu.co/ (last accessed: January 15, 2016) | 1 | 1 |
| BGCI (2017). GlobalTreeSearch online database, Available at: www.bgci.org/globaltree_search.php (last accessed: August 14, 2017) | 1 | 1 |
| Bingham, M.G., Willemen, A., Wursten, B.T., Ballings, P. & Hyde, M.A. (2016). Flora of Zambia, Available at: http://www.zambiaflora.com/ (last accessed: November 14, 2016) | 1 | 0 |
| BioScripts (2014). Flora Vascular, Available at: http://www.floravascular.com/ (last accessed: May 25, 2014) | 1 | 1 |
| Borges, P.A.V., Abreu, C., Aguiar, A.M.F., Carvalho, P., Jardim, R., Melo, I., Oliveira, P., Sérgio, C., Serrano, A.R.M. & Vieira, P. (2008). *Listagem dos fungos, flora e fauna terrestres dos arquipélagos da Madeira e Selvagens*. Direcção Regional do Ambiente da Madeira and Universidade dos Açores, Funchal and Angra do Heroísmo, Portugal | 1 | 0 |
| Borges, P.A.V., Costa, A., Cunha, R., Gabriel, R., Gonçalves, V., Martins, A.F., Melo, I., Parente, M., Raposeiro, P., Rodrigues, P., Santos, R.S., Silva, L., Vieira, P. & Vieira, V. (2010). *A list of the terrestrial and marine biota from the Azores*. Princípiã, Cascais | 1 | 0 |
| Botanical Garden Tel Aviv (2015). Flora of Isreal (Pers. Comm.) | 1 | 1 |
| Bowdoin Scientific Station (2011). Vascular plants of Kent Island, Available at: https://www.bowdoin.edu/kent-island/species/plants.shtml (last accessed: September 14, 2011) | 1 | 0 |
| Breckle, S.-W., Hedge, I.C. & Rafiqpoor, M.D. (2013). *Vascular plants of Afghanistan. An augmented checklist*. Scientia Bonnensis, Bonn | 1 | 0 |
| Brennan, K. (1996). *An annotated checklist of the vascular plants of the Alligator Rivers Region, Northern Territory, Australia*. Supervising Scientist, Barton, Australia | 1 | 1 |
| Bridgewater, S.G.M., Harris, D.J., Whitefoord, C., Monro, A.K., Penn, M.G., Sutton, D.A., Sayer, B., Adams, B., Balick, M.J., Atha, D.H., Solomon, J. & Holst, B.K. (2006). A Preliminary Checklist of the vascular plants of the Chiquibul Forest, Belize. *Edinburgh Journal of Botany*, **63**, 269–321 | 0 | 1 |
| Brofas, G., Karetos, G., Panitsa, M. & Theocharopoulos, M. (2001). The flora and vegetation of Gyalí Island, SE Aegean, Greece. *Willdenowia*, **31**, 51–70 | 1 | 0 |
| Broughton, D.A. & McAdam, J.H. (2005). A checklist of the native vascular flora of the Falkland Islands (Islas Malvinas). new information on the species present, their ecology, status and distribution. *The Journal of the Torrey Botanical Society*, **132**, 115–148 | 1 | 0 |
| Brundu, G. & Camarda, I. (2013). The Flora of Chad: a checklist and brief analysis. *Phytokeys*, **23**, 1–17 | 1 | 0 |
| Bundesamt für Naturschutz (2016). Floraweb, Available at: www.floraweb.de (last accessed: September 15, 2016) | 1 | 1 |
| Burton, R.M. (1991). A check-list and evaluation of the flora of Nisyros (Dodecanese, Greece). *Willdenowia*, **20**, 15–38 | 1 | 0 |
| Butler, B.J., Barclay, J.S. & Fisher, J.P. (1999). Plant communities and flora of Robins Island (Long Island), New York. *Journal of the Torrey Botanical Society*, **126**, 63–76 | 1 | 0 |
| Byrd, G.V. (1984). Vascular vegetation of Buldir Island, Aleutian Islands, Alaska, compared to another Aleutian Island. *Arctic*, **37**, 37–48 | 1 | 0 |
| CARMABI (2009). Dutch Caribbean Biodiversity Explorer, Available at: http://www.dcbiodata.net/explorer/home (last accessed: June 24, 2011) | 1 | 0 |
| Cascante-Marín, A. & Estrada-Chavarría, A. (2012). Las plantas vasculares de El Rodeo, Costa Rica. *Brenesia*, **77**, 71–128 | 1 | 1 |
| Case, T.J., Cody, M.L. & Ezcurra, E. (2002). *A new island biogeography of the Sea of Cortés*. Oxford University Press, New York, NY | 1 | 0 |
| Catarino, L., Martins, E.S., Basto, M.F. & Diniz, M.A. (2008). An annotated checklist of the vascular flora of Guinea-Bissau (West Africa). *Blumea-Biodiversity, Evolution and Biogeography of Plants*, **53**, 1–222 | 1 | 0 |
| Chang, C.-S., Kim, H. & Chang, K. *Provisional Checklist of the Vascular Plants for the Korea Peninsular Flora (KPF). Version 1.0*, Korea | 1 | 0 |
| Chawla, A., Parkash, O., Sharma, V., Rajkumar, S., Lal, B., Gopichand, Singh, R.D. & Thukral, A.K. (2012). Vascular plants, Kinnaur, Himachal Pradesh, India. *Check list*, **8**, 321–348 | 0 | 1 |
| Cheffings, C.M. & Farrell, L. (eds.) (2005). *The vascular plant red data list for Great Britain*. Joint Nature Conservation Committee, Peterborough | 1 | 0 |
| Chernyaeva, A.M. (1973). Flora of Onkotan Island. *Bulletin of Main Botanical Garden*, **87**, 21–29 | 1 | 0 |
| Chiapella, J. & Ezcurra, C. (1999). La flora del parque provincial Tromen, provincia de Neuquén, Argentina. *Multequina*, **8**, 51–60 | 1 | 0 |
| Chinese Virtual Herbarium (2016). The Flora of China v. 5.0, Available at: http://www.cvh.org.cn/ (last accessed: January 15, 2016) | 1 | 1 |
| Chong, K.Y., Tan, T.W.H. & Corlett, R.T. (2009). *A checklist of the total vascular plant flora of Singapore. Native, Naturalised and Cultivated Species*. Raffles Museum of Biodiversity Research, Singapore | 1 | 1 |
| Christmas Island National Park (2002). *Third Christmas Island national park management plan*. Parks Australia North, Christmas Island, Australia | 1 | 0 |
| Christodoulakis, D. (1996). The flora of Ikaria (Greece, E. Aegean Islands). *Phyton*, **36**, 63–91 | 1 | 0 |
| Clark, J.L., Neill, D.A. & Asanza, M. (2006). Floristic checklist of the Mache-Chindul mountains of Northwestern Ecuador. *Contributions from the United States National Herbarium*, **54**, 1–180 | 1 | 1 |
| Cochard, R. & Bloesch, U. (2007). Electronic plant species database of the Saadani National Park, coastal Tanzania, Available at: http://www.wildlife-baldus.com/saadani.html (last accessed: November 14, 2016) | 1 | 1 |
| CONABIO (2016). Sistema Nacional de Información sobre Biodiversidad, Available at: https://www.gob.mx/conabio (last accessed: April 11, 2016) | 1 | 0 |
| Conti, F. & Bartolucci, F. (2015). *The Vascular Flora of the National Park of Abruzzo, Lazio and Molise (Central Italy)*. Springer International Publishing, Cham | 1 | 0 |
| Conti, F., Abbate, G., Alessandrini, A. & Blasi, C. (2005). *Annotated Checklist of the Italian Vascular Flora*. Palombi Editori, Roma, Italy | 1 | 0 |
| Convey, P., Lewis Smith, R.I., Hodgson, D.A. & Peat, H.J. (2000). The flora of the South Sandwich Islands, with particular reference to the influence of geothermal heating. *Journal of Biogeography*, **27**, 1279–1295 | 1 | 0 |
| Costion, C. & Lorence, D. (2012). The endemic plants of Micronesia. A geographical checklist and commentary. *Micronesica*, **43**, 51–100 | 1 | 1 |
| Cronk, Q.C.B. (1989). The past and present vegetation of St Helena. *Journal of Biogeography*, **16**, 47–64 | 1 | 0 |
| Da Vela, M., Frignani, F., Bonari, G. & Angiolini, C. (2013). La flora vascolare della riserva naturale "La Pietra" (Toscana meridionale). *Micologia e vegetazione mediterranea*, **28**, 135–160 | 0 | 1 |
| Danihelka, J., Chrtek, J. & Kaplan, Z. (2012). Checklist of vascular plants of the Czech Republic. *Preslia*, **84**, 647–811 | 1 | 0 |
| D'Arcy, W.G. (1971). The island of Anegada and its flora. *Atoll Research Bulletin*, **139**, 1–21 | 1 | 0 |
| Dauby, G., Leal, M. & Stevart, T. (2008). Vascular plant checklist of the coastal National Park of Pongara, Gabon. *Systematics and geography of plants*, **78**, 155–216 | 0 | 1 |
| Dauby, G., Zaiss, R., Blach-Overgaard, A., et al. (2016). RAINBIO. A mega-database of tropical African vascular plants distributions. *PhytoKeys*, **74**, 1–18 | 1 | 1 |
| de la Luz, León, Rebman, J., Domínguez-León, M. & Domínguez-Cadena, R. (2008). The vascular flora and floristic relationships of the sierra de la Giganta in Baja California Sur, México. *Revista mexicana de biodiversidad*, **79**, 29–65 | 0 | 1 |
| Desmet, P. & Brouillet, L. (2013). Database of Vascular Plants of Canada (VASCAN): a community contributed taxonomic checklist of all vascular plants of Canada, Saint Pierre and Miquelon, and Greenland. *Phytokeys*, **25**, 55–67 | 0 | 1 |
| Dimopoulos, P., Raus, T., Bergmeier, E., Constantinidis, T., Iatrou, G., Kokkini, S., Strid, A. & Tzanoudakis, D. (2013). *Vascular plants of Greece. An annotated checklist*. Botanischer Garten und Botanisches Museum Berlin-Dahlem, Freie Universität Berlin; Hellenic Botanical Society, Berlin, Athens | 1 | 0 |
| Domínguez, E., Marticorena, C., Elvebakk, A. & Pauchard, A. (2004). Catálogo de la flora vascular del Parque Nacional Pali Aike. XII Región, Chile. *Gayana Botánica*, **61**, 67–72 | 1 | 1 |
| Doroftei, M., Oprea, A., Ștefan, N. & Sârbu, I. (2011). Vascular wild flora of Danube Delta Biosphere Reserve. *Sci. Annals of Danube Delta Institute*, **17**, 15–52 | 1 | 0 |
| Dowhan, J.J. & Rozsa, R. (1989). Flora of Fire Island, Suffolk County, New York. *Bulletin of the Torrey Botanical Club*, **116**, 265–282 | 1 | 0 |
| Du Puy, D.J. (1993). Christmas Island. species lists, Available at: http://www.anbg.gov.au/abrs/online-resources/flora/ (last accessed: April 06, 2011) | 1 | 0 |
| Egea, J. de, Peña-Chocarro, M., Espada, C. & Knapp, S. (2012). Checklist of vascular plants of the Department of Ñeembucú, Paraguay. *Phytokeys*, **9**, 15–179 | 1 | 1 |
| Egorova, E.M. (1964). Flora of Shishkotan Island. *Bulletin of the Main Botanical Garden*, **54**, 114–120 | 1 | 0 |
| Ejaz-ul-Islam Dar, M., Cochard, R., Shrestha, R.P. & Ahmad, S. (2012). Floristic composition of Machiara National Park, District Muzaffarabad Azad Kashmir, Pakistan. *International Journal of Biosciences*, **2**, 28–45 | 0 | 1 |
| Engemann, K., Sandel, B., Boyle, B.L., Enquist, B.J., Jørgensen, P.M., Kattge, J., McGill, B.J., Morueta-Holme, N., Peet, R.K., Spencer, N.J., Violle, C., Wiser, S.K. & Svenning, J.-C. (2016). A plant growth form dataset for the New World. *Ecology*, **97**, 3243 | 0 | 1 |
| Esler, A.E. (1978). Botanical features of the Mokohinau Islands. *TANE*, **24**, 187–197 | 1 | 0 |
| Espinosa-Jiménez, J.A., Pérez-Farrera, M.Á. & Martínez-Camilo, R. (2011). Inventario florístico del Parque Nacional Cañón del Sumidero, Chiapas, México. *Boletín de la Sociedad Botánica de México*, **89**, 37–82 | 0 | 1 |
| Evarts-Bunders, P., Evarte-Bundere, G., Băra, J. & Nitcis, M. (2013). The flora of vascular plants in nature reserve „Eglone”. *Acta Biologica Universitatis Daugavpiliensis*, **13**, 21–38 | 1 | 0 |
| Exell, A.W. (1944). *Catalogue of the vascular plants of S. Tome (with Principe and Annobon)*. Trustees of the British Museum, London, UK | 1 | 0 |
| Fern, K. & Fern, A. (2015). Useful Tropical Plants Database (last accessed: July 25, 2015) | 0 | 1 |
| Ferreira, A.L., Coutinho, B.R., Pinheiro, H.T. & Thomaz, L.D. (2007). Composição florística e formações vegetais da Ilha dos Franceses, Espírito Santo. *Bol. Mus. Biol. Mello Leitao*, **22**, 25–44 | 0 | 1 |
| Figueiredo, E., Paiva, J., Stevart, T., Oliveira, F. & Smith, G.F. (2011). Annotated catalogue of the flowering plants of São Tomé and Príncipe. *Bothalia*, **41**, 41–82 | 1 | 1 |
| Figueroa-C., Y. & Galeano, G. (2007). Lista comentada de las plantas vasculares del enclave seco interandino de La Tatacoa (Huila, Colombia). *Caldasia*, **29**, 263–281 | 1 | 1 |
| Fischer, E., Rembold, K., Althof, A., Obholzer, J., Malombe, I., Mwachala, G., Onyango, J.C., Dumbo, B. & Theisen, I. (2010). Annotated checklist of the vascular plants of Kakamega Forest, Western Province, Kenya. *Journal of East African Natural History*, **99**, 129–226 | 1 | 1 |
| Fischer, M.A., Adler, W. & Oswald, K. (2008). *Exkursionsflora für Österreich, Liechtenstein und Südtirol. Bestimmungsbuch für alle in der Republik Österreich, im Fürstentum Liechtenstein und in der Autonomen Provinz Bozen, 3., verb. u. erw. Aufl. der "Exkursionsflora von Österreich"*. OÖ Landesmuseum, Linz | 1 | 0 |
| Florence, J., Chevillotte, H., Ollier, C. & Meyer, J.-Y. (2007). Base de données botaniques Nadeaud de l'Herbier de la Polynésie française (PAP), Available at: http://www.herbier-tahiti.pf/ (last accessed: July 01, 2011) | 1 | 0 |
| Fosberg, F.R., Renvoize, S.A. & Townsend, C.C. (1980). *The flora of Aldabra and neighbouring islands*. HMSO, London, UK | 1 | 0 |
| Franklin, J., Keppel, G. & Whistler, W.A. (2008). The vegetation and flora of Lakeba, Nayau and Aiwa islands, central Lau Group, Fiji. *Micronesica*, **40**, 169–225 | 1 | 0 |
| Funk, V.A., Hollowell, T., Berry, P., Kelloff, C. & Alexander, S.N. (2007). *Checklist of the plants of the Guiana Shield (Venezuela: Amazonas, Bolivar, Delta Amacuro; Guyana, Surinam, French Guiana)*. Department of Botany, National Museum of Natural History, Washington, DC | 1 | 0 |
| Gabrielsen, G.W., Brekke, B., Alsos, I.G. & Hansen, J.R. (1997). *Natur-og kulturmiljøet på Jan Mayen*. Norsk Polarinstitut, Oslo, Norway | 1 | 0 |
| Gage, S., Joneson, S.L., Barkalov, V.Y., Eremenko, N.A. & Takahashi, H. (2006). A newly compiled checklist of the vascular plants of the Habomais, the Little Kurils. *Bulletin of the Hokkaido University Museum*, **3**, 67–91 | 1 | 0 |
| Gamit, S.B., Maurya, R.R., Qureshi, U.M. & Solanki, H.A. (2015). Check list of flowering plants in Tapi District, Gujarat, India. *International Journal of advanced Research*, **3**, 1104–1123 | 0 | 1 |
| Gerlach, J. (2003). The biodiversity of the granitic islands of Seychelles. *Phelsuma*, **11** (Supplement A), 1–47 | 1 | 0 |
| Gray, A., Pelembe, T. & Stroud, S. (2005). The conservation of the endemic vascular flora of Ascension Island and threats from alien species. *Oryx*, **39**, 449–453 | 1 | 0 |
| Green, P.S. (1994). Lord Howe Island. species lists, Available at: http://www.environment.gov.au/biodiversity/abrs/online-resources/flora/49/index.html (last accessed: April 06, 2011) | 1 | 0 |
| Greene, S.W. & Walton, D.W.H. (1975). An annotated check list of the sub-Antarctic and Antarctic vascular flora. *Polar Record*, **17**, 473–484 | 1 | 0 |
| Harris, D.J. (2002). *The vascular plants of the Dzanga-Sangha Reserve, Central African Republic*. Royal Botanic Garden Edinburgh, Edinburgh, Scotland, UK | 1 | 1 |
| Hauenstein, E., Muñoz-Pedreros, A., Yáñez, J., Sánchez, P., Möller, P., Guiñez, B. & Gil, C. (2009). Flora y vegetación de la Reserva Nacional Lago Peñuelas, Reserva de la Biosfera, Región de Valparaíso, Chile. *Bosque (Valdivia)*, **30**, 159–179 | 1 | 0 |
| Hawkins, B.A., Rueda, M., Rangel, T.F., Field, R., Diniz-Filho, J.A.F. & Linder, P. (2013). Community phylogenetics at the biogeographical scale: cold tolerance, niche conservatism and the structure of North American forests. *Journal of Biogeography*, **41**, 23–38 | 0 | 1 |
| Heberling, J.M., Jo, I., Kozhevnikov, A., Lee, H. & Fridley, J.D. (2017). Biotic interchange in the Anthropocene. Strong asymmetry in East Asian and eastern North American plant invasions. *Global Ecology and Biogeography*, **26**, 447–458 | 0 | 1 |
| Hill, M.J. (ed.) (2002). *Biodiversity surveys and conservation potential of inner Seychelles islands*. Smithsonian Institution, Washington, DC | 1 | 0 |
| Hill, S.R. (1986). An annotated checklist of the vascular flora of Assateague Island (Maryland and Virginia). *Castanea*, **51**, 265–305 | 1 | 0 |
| Hnatiuk, R.J. (1993). Subantarctic Islands. species lists, Available at: http://www.environment.gov.au/biodiversity/abrs/online-resources/flora/50/index.html (last accessed: April 07, 2011) | 1 | 0 |
| Hyde, M.A., Wursten, B.T., Ballings, P. & Coates Palgrave, M. (2016). Flora of Botswana, Available at: http://www.botswanaflora.com/ (last accessed: November 14, 2016) | 1 | 0 |
| Hyde, M.A., Wursten, B.T., Ballings, P. & Coates Palgrave, M. (2016). Flora of Zimbabwe (last accessed: November 14, 2016) | 1 | 0 |
| Hyde, M.A., Wursten, B.T., Ballings, P. & Coates Palgrave, M. (2016). Flora of Mozambique, Available at: http://www.mozambiqueflora.com/ (last accessed: November 14, 2016) | 1 | 0 |
| Hyde, M.A., Wursten, B.T., Ballings, P. & Coates Palgrave, M. (2016). Flora of Malawi, Available at: http://www.malawiflora.com/index.php (last accessed: November 14, 2016) | 1 | 0 |
| Imada, C.T. (ed.) (2012). *Hawaiian Native and Naturalized Vascular Plants Checklist*, (December 2012 update). Hawaii Biological Survey, Bishop Museum, Honolulu, Hawai'i | 1 | 0 |
| INBIO (2000). Lista de planta de Costa Rica. With updates by Eduardo Chacón, Available at: http://www.inbio.ac.cr/papers/manual_plantas/index.html (last accessed: February 05, 2016) | 1 | 0 |
| Jacks, B.R. (2010). *Plants of Magnetic Island*, 3rd edn. James Cook University, Townsville, Australia | 1 | 0 |
| Jahn, R. & Schönfelder, P. (1995). *Exkursionsflora für Kreta*. Ulmer (Eugen), Stuttgart, Germany | 1 | 0 |
| Jaramillo Díaz, P. & Guézou, A. (2011). CDF checklist of Galapagos vascular plants. Bungartz, F., Herrera, H.W., Jaramillo, P., Tirado, N., Jiménez-Uzategui, G., Ruiz, D., Guézou, A. & Ziemmeck, F. (eds.), Available at: http://www.darwinfoundation.org/datazone/checklists/vascular-plants/ (last accessed: February 02, 2011) | 1 | 0 |
| Jardim Botânico do Rio de Janeiro (2016). Flora do Brasil 2020 em construção, Available at: http://floradobrasil.jbrj.gov.br/ (last accessed: May 09, 2016) | 1 | 1 |
| Jiménez, J.E., Juárez, P. & Díaz, A. (2016). Checklist of the vascular flora of Reserva Biológica San Luis, Costa Rica. *Check list*, **12**, 1859 | 1 | 1 |
| Johnson, P.N. & Campbell, D.J. (1975). Vascular plants of the Auckland Islands. *New Zealand journal of botany*, **13**, 665–720 | 1 | 0 |
| Johnston, I.M. (1931). The flora of the Revillagigedo Islands. *Proceedings of the California Academy of Sciences*, **20**, 9–104 | 1 | 0 |
| Jordano, P. (2008). FRUBASE, Available at: http://ebd10.ebd.csic.es/mywork/frubase/frubase.html (last accessed: 26.06.15) | 0 | 1 |
| Junak, S., Philbrick, R., Chaney, S. & Clark, R. (1997). *A checklist of vascular plants of Channel Islands National Park*, 2nd edn. Southwest Parks and Monuments Association, Tucson, Arizona | 1 | 0 |
| Kabuye, C.H.S., Mungai, G.M. & Mutangah, J.G. (1986). Flora of Kora National Reserve. *Kora: An Ecological Inventory of the Kora National Reserve, Kenya. Kora Research Project 1982-85 : a Joint Venture Between the National Museums of Kenya and the Royal Geographical Society* (ed. by M. Coe and N.M. Collins), pp. 57–104. Royal Geographical Society | 0 | 1 |
| Kamari, G., Phitos, D., Snogerup, B. & Snogerup, S. (1988). Flora and vegetation of Yioura, N Sporades, Greece. *Willdenowia*, **17**, 59–85 | 1 | 0 |
| Karlsson, T. & Agestam, M. (2014). Checklist of Nordic vascular plants. Sweden Checklist, Available at: http://www.euphrasia.nu/checklista/index.eng.html (last accessed: February 11, 2016) | 1 | 0 |
| Kattge, J., Diaz, S., Lavorel, S., Prentice, I.C., Leadley, P., Bönsch, G., Garnier, E., Westoby, M., Reich, P.B. & Wright, I.J. (2011). TRY – a global database of plant traits. *Global Change Biology*, **17**, 2905–2935 | 0 | 1 |
| Kelloff, C.L. & Funk, V.A. (1998). *Preliminary checklist of the plants of Kaieteur National Park, Guyana*. National Museum of Natural History, Smithsonian Institution, Washington | 1 | 0 |
| Kelly, L. (2006). The vascular flora of Huggins Island, Onslow County, North Carolina. *Castanea*, **71**, 295–311 | 1 | 0 |
| Kemenes, A. (2003). Distribuição espacial da flora terrestre fanerogâmica do Parque Nacional Marinho de Abrolhos, BA. *Revista Brasil. Bot*, **26**, 141–150 | 0 | 1 |
| Keppel, G. (2005). *Summary report on forests of the Mataqali Nadicake Kilaka, Kubulau district, Bua, Vanua Levu* | 0 | 1 |
| Keppel, G., Gillespie, T.W., Ormerod, P. & Fricker, G.A. (2016). Habitat diversity predicts orchid diversity in the tropical south-west Pacific. *Journal of Biogeography*, **43**, 2332–2342 | 1 | 0 |
| Kerguelen, M. (2005). Base de Données Nomenclaturales de la Flore de France, Available at: http://www.tela-botanica.org/page:284# (last accessed: January 13, 2012) | 1 | 0 |
| Khanina, L., Zaugolnova, L., Smirnova, O., Shovkun, M. & Glukhova, E. (2016). Flora of vascular plants in the Central European Russia, Available at: http://www.impb.ru/eco/index.php?l=en (last accessed: September 06, 2017) | 1 | 0 |
| Kim, H.-J., Ji, S.-J., Jung, S.-Y., Park, S.H., Lee, S.-G., Lee, C.-W. & Chang, K.S. (2015). Flora of Vascular Plants in Deokjeokdo (Ongjin-gun) and Its Adjacent Regions, Korea. *Korean Journal of Plant Resources*, **28**, 487–510 | 1 | 0 |
| Kingston, N., Waldren, S. & Bradley, U. (2003). The phytogeographical affinities of the Pitcairn Islands – a model for south-eastern Polynesia? *Journal of Biogeography*, **30**, 1311–1328 | 1 | 1 |
| Kirchner, F., Picot, F., Merceron, E. & Gigot, G. (2010). *Flore vasculaire de La Réunion*. Conservatoire Botanique National de Mascarin, Réunion, France | 1 | 0 |
| Kleyer, M., Bekker, R.M., Knevel, I.C., et al. (2008). The LEDA Traitbase: a database of life-history traits of the Northwest European flora. *Journal of Ecology*, **96**, 1266–1274 | 0 | 1 |
| Koltzenburg, M. (2011). Checkliste der Gefässpflanzen Irlands, Available at: http://www.saxifraga.de/eire/irl_artenlisten_gp.html (last accessed: April 04, 2011) | 1 | 0 |
| Kraaij, T. (2011). The flora of the Bontebok National Park in regional perspective. *South African Journal of Botany*, **77**, 455–473 | 1 | 1 |
| Kraft, T.S., Wright, S.J., Turner, I., Lucas, P.W., Oufiero, C.E., Supardi Noor, M.N., Sun, I.-F., Dominy, N.J. & Whitney, K. (2015). Seed size and the evolution of leaf defences. *Journal of Ecology*, **103**, 1057–1068 | 0 | 1 |
| Kristinsson, H. (2008). *Checklist of the vascular plants of Iceland*. Náttúrufræðistofnun Íslands, Reykjavík, Iceland | 1 | 0 |
| Kumar, A., Bajpai, O., Mishra, A.K., Sahu, N., Behera, S.K., Bargali, S.S. & Chaudhary, L.B. (2015). A checklist of the flowering plants of Katarniaghat Wildlife Sanctuary, Uttar Pradesh, India. *Journal of Threatened Taxa*, **7**, 7309–7408 | 0 | 1 |
| Kuzmenkova, S.M. (2015). Plants of Belarus, Available at: http://hbc.bas-net.by/plantae/eng/default.php (last accessed: February 15, 2016) | 1 | 0 |
| Lange, P.J. de & Cameron, E.K. (1999). The vascular flora of Aorangi Island, Poor Knights Islands, northern New Zealand. *New Zealand journal of botany*, **37**, 433–468 | 1 | 0 |
| Lange, P.J. de, Heenan, P.B. & Rolfe, J.R. (2011). *Checklist of vascular plants recorded from Chatham Islands* | 1 | 0 |
| Lazkov, G.A. & Sultanova, B.A. (2011). *Checklist of vascular plants of Kyrgyzstan*. Botanical Museum, Finnish Museum of Natural History, Helsinki | 1 | 0 |
| Le Houerou, H.N. (2004). Plant diversity in Marmarica (Libya & Egypt): a catalogue of the vascular plants reported with their biology, distribution, frequency, usage, | 1 | 1 |
| economic potential, habitat and main ecological features, with an extensive bibliography. *Candollea*, **59**, 259–308 |  |  |
| Lee, R.-Y., Jang, H.-D., Kim, Y.-Y., Yang, S.-G., Choi, H.-J., Ji, S.-J. & Oh, B.-U. (2014). Flora of vascular plants in the Chilgapsan Provincial Park, Korea. *Journal of Asia-Pacific Biodiversity*, **7**, 237–247 | 1 | 0 |
| Lester-Garland, L.V. (1903). *A flora of the islands of Jersey. with a list of the plants of the Channel Islands in general, and remarks upon their distribution and geographical affinities*. West, Newman & Co, London, UK | 1 | 0 |
| Letcher, S.G., Lasky, J.R., Chazdon, R.L., et al. (2015). Environmental gradients and the evolution of successional habitat specialization. A test case with 14 Neotropical forest sites. *Journal of Ecology*, **103**, 1276–1290 | 0 | 1 |
| Levin, G.A. & Moran, R. (1989). The vascular flora of Socorro, Mexico. *Memoirs of the San Diego Society of Natural History*, **16**, 1–71 | 1 | 0 |
| Limbu, D., Koirala, M. & Shang, Z. (2013). A Checklist of Angiospermic Flora of Tinjure-Milke-Jaljale, Eastern Nepal. *Nepal Journal of Science and Technology*, **13**, 87–96 | 0 | 1 |
| Linhart, Y.B. (1980). Local biogeography of plants on a Caribbean atoll. *Journal of Biogeography*, **7**, 159–171 | 0 | 1 |
| Lipkin, R. (2005). *Aniakchak National Monument and Preserve, vascular plant inventory: final technical report*. National Park Service, Southwest Alaska Network Inventory & Monitoring Program, Anchorage, USA | 1 | 0 |
| Lopez-Martinez, J.O., Sanaphre-Villanueva, L., Dupuy, J.M., Hernandez-Stefanoni, J.L., Meave, J.A. & Gallardo-Cruz, J.A. (2013). Beta-Diversity of functional groups of woody plants in a tropical dry forest in Yucatan. *PloS one*, **8**, e73660 | 0 | 1 |
| Lord, J.M. (2015). Patterns in floral traits and plant breeding systems on Southern Ocean Islands. *AoB Plants*, **7** | 1 | 1 |
| Lorite, J. (2016). An updated checklist of the vascular flora of Sierra Nevada (SE Spain). *Phytotaxa*, **261**, 1–57 | 1 | 0 |
| Luke, Q. (2005). Annotated Checklist of the Plants of the Shimba Hills, Kwale District, Kenya. *Journal of East African Natural History*, **94**, 5–120 | 1 | 1 |
| Luna-Jorquera, G., Fernández, C.E. & Rivadeneira, M.M. (2012). Determinants of the diversity of plants, birds and mammals of coastal islands of the Humboldt current systems. Implications for conservation. *Biodiversity and Conservation*, **21**, 13–32 | 1 | 0 |
| Marquand, E.D. (1901). *Flora of Guernsey and the lesser Channel Islands. namely Alderney, Sark, Herm, Jethou, and the adjacent islets*. Dulau & Co, London, UK | 1 | 0 |
| Marticorena, C., Squeo, F.A., Arancio, G. & Muñoz, M. (2008). Catálogo de la flora vascular de la IV Región de Coquimbo. *Libro rojo de la flora nativa y de los sitios prioritarios para su conservación: Región de Atacama* (ed. by F.A. Squeo, G. Arancio and J.R. Gutiérrez), pp. 105–142. Ediciones Universidad de La Serena La Serena | 1 | 1 |
| Marticorena, C., Stuessy, T.F. & Baeza, C.M. (1998). Catalogue of the vascular flora of the Robinson Crusoe or Juan Fernández islands, Chile. *Gayana Botánica*, **55**, 187–211 | 1 | 1 |
| Masharabu, T. *Flore et végétation du Parc National de la Ruvubu au Burundi: diversité, structure et implications pour la conservation*. Editions universitaires europeennes | 1 | 1 |
| McClatchey, W., Thaman, R. & Vodonaivalu, S. (2000). A preliminary checklist of the flora of Rotuma with Rotuman names. *Pacific Science*, **54**, 345–363 | 1 | 0 |
| McCrea, J. (2003). *Inventory of the land conservation values of the Houtman Abrolhos Islands*. Department of Fisheries, Government of Western Australia, Perth, Australia | 1 | 0 |
| Medina, R., Reina-E, M., Herrera, E., Ávila, F.A., Chaparro, O. & Cortés-B., R. (2010). Catálogo preliminar da flora vascular dos bosques subandinos da cuchilla El Fara (Santander-Colômbia). *Colombia Forestal*, **13**, 55–85 | 0 | 1 |
| Medjahdi, B., Ibn Tattou, M., Barkat, D. & Benabedli, K. (2009). La flore vasculaire des Monts des Trara (Nord Ouest Algérie). *Acta Botanica Malacitana*, **34**, 57–75 | 1 | 0 |
| Miller, A.G. & Morris, M. (2004). *Ethnoflora of the Soqatra Archipelago*. Royal Botanic Garden, Edinburgh, UK | 1 | 0 |
| Ministry of Environment (2012). *The national red list 2012 of Sri Lanka. Conservation status of the fauna and flora*, Colombo, Sri Lanka | 1 | 0 |
| Moran, R. (1996). *The flora of Guadalupe Island, Mexico*. California Academy of Sciences, San Francisco, CA | 1 | 0 |
| Morat, P., Jaffré, T., Tronchet, F., Munziger, J., Pillon, Y., Veillon, J.-M., Chalopin, M., Birnbaum, P., Rigault, F., Dagostini, G., Tinerl, J. & Lowry II, P.P. (2012). The taxonomic reference base Florical and characteristics of the native vascular flora of New Caledonia. *Adansonia*, **34**, 179–221 | 1 | 0 |
| Nationalpark Eifel (2015). Artenliste Farne und Blütenpflanzen, Available at: http://www.nationalpark-eifel.de/go/artenliste.html (last accessed: March 11, 2015) | 1 | 0 |
| Nikolić, T. (2016). Flora Croatica Database, Available at: http://hirc.botanic.hr/fcd (last accessed: February 12, 2016) | 1 | 0 |
| Norton, J., Majid, S.A., Allan, D., Al Safran, M., Böer, B. & Richer, R.A. (2009). *An illustrated checklist of the flora of Qatar*. Browndown Publications Gosport, Gosport, UK | 1 | 1 |
| Notov, A.A. (2010). *National park "Zavidovo": vascular plants, bryophyte, lichens*, Moscow | 1 | 0 |
| NPS (2015). NPSpecies. Information on Species in National Parks, Available at: https://irma.nps.gov/NPSpecies/ (last accessed: April 09, 2015) | 1 | 0 |
| Oggero, A.J. & Arana, M.D. (2012). Inventario de las plantas vasculares del sur de la zona serrana de Córdoba, Argentina. *Hoehnea*, **39**, 171–199 | 1 | 0 |
| Oliveira-Filho, A.T. (2014). NeoTropTree. Flora arbórea da Região Neotropical: Um banco de dados envolvendo biogeografia, diversidade e conservação, Available at: http://www.icb.ufmg.br/treetatlan/ (last accessed: February 12, 2016) | 0 | 1 |
| Pal, D., Kumar, A. & Dutt, B. (2014). Floristic diversity of Theog Forest Division, Himachal Pradesh, Western Himalaya. *Check list*, **10**, 1083–1103 | 1 | 1 |
| Pandža, M. & Skvorc, Z. (2002). The flora of some uninhabited Sibenik Archipelago islands (Dalmatia, Croatia). *Natura Croatica*, **11**, 367–385 | 1 | 0 |
| Pandža, M. (2002). Flora of the small islands of Murter. *Natura Croatica*, **11**, 77–101 | 1 | 0 |
| Pandža, M. (2003). Flora of the island of Zirje and the small islands around it (eastern Adriatic coast, Croatia). *Acta Botica Croatia*, **62**, 115–139 | 1 | 0 |
| Pandža, M. (2010). Flora parka prirode Papuk (Slavonija, Hrvatska). *Šumarski list*, **134**, 25–43 | 1 | 0 |
| Parks Canada (2015). Biotics Web Explorer, Available at: http://www.pc.gc.ca/apps/bos/BOSIntro_e.asp (last accessed: April 13, 2015) | 1 | 0 |
| Paula, S., Arianoutsou, M., Kazanis, D., Tavsanoğlu, Ç., Lloret, F., Buhk, C., Ojeda, F., Luna, B., Moreno, J.M. & Rodrigo, A. (2009). Fire-related traits for plant species of the Mediterranean Basin. *Ecology*, **90**, 1420 | 0 | 1 |
| Peña-Chocarro, M.d.C. (2010). *Updated checklist of vascular plants of the Mbaracayú Forest Nature Reserve (Reserva Natural del Bosque Mbaracayú), Paraguay*. Magnolia Press, Auckland, N.Z. | 1 | 1 |
| Pôle Flore Habitats (2015). Catalogue de la flore vasculaire de Rhône-Alpes, Available at: http://www.pifh.fr/pifhcms/index.php (last accessed: January 12, 2015) | 1 | 0 |
| Price, J.P. & Wagner, W.L. (2011). A phylogenetic basis for species–area relationships among three Pacific Island floras. *American Journal of Botany*, **98**, 449–459 | 1 | 0 |
| Proctor, G.R. (1980). Checklist of the plants of Little Cayman. Geography and ecology of Little Cayman. *Atoll Research Bulletin*, **241**, 71–80 | 1 | 0 |
| Proctor, G.R. (1989). *Ferns of Puerto Rico and the Virgin Islands*. Memoirs of the New York Botanical Garden, New York, NY | 1 | 0 |
| Queensland Government (2014). Census of the Queensland flora 2014, Available at: https://data.qld.gov.au/dataset/census-of-the-queensland-flora-2014 (last accessed: February 05, 2015) | 1 | 0 |
| Rahman, A.H.M.M. (2013). Angiospermic Flora of Rajshahi District, Bangladesh. *American Journal of Life Sciences*, **1**, 105 | 0 | 1 |
| Rahman, M.S., Hossain, G.M., Khan, S.A. & Uddin, S.N. (2015). An annotated Checklist of the Vascular Plants of Sundarban Mangrove Forest of Bangladesh. *Bangladesh Journal of Plant Taxonomy*, **22**, 17–41 | 0 | 1 |
| Rakov, N.S., Saksonov S.V., Senator S.A. & Vasjukov V.M. (2014). *Vascular plants of Ulyanovsk Region*. Russian Academy of Sciences, Togliatti | 1 | 0 |
| Raulerson, L. (2006). Checklist of plants of the Mariana Islands. *University of Guam Herbarium Contribution*, **37**, 1–69 | 1 | 0 |
| Renvoize, S.A. (1975). A floristic analysis of the western Indian Ocean coral islands. *Kew Bulletin*, **30**, 133–152 | 1 | 0 |
| Robinson, A.C., Canty, P.D. & Fotheringham, D. (2008). Investigator group expedition 2006. flora and vegetation. *Transactions of the Royal Society of South Australia*, **132**, 173–220 | 1 | 0 |
| Robinson, A.C., Canty, P.D., Wace, N.M. & Barker, R.M. (2003). The encounter 2002 expedition to the isles of St Francis, South Australia. flora and vegetation. *Transactions of the Royal Society of South Australia*, **127**, 107–128 | 1 | 0 |
| Rossetto, E.F.S. & Vieira, A.O.S. (2013). Vascular Flora of the Mata dos Godoy State Park, Londrina, Paraná, Brazil. *Check list*, **9**, 1020–1034 | 1 | 1 |
| Roux, J.P. (2009). *Synopsis of the Lycopodiophyta and Pteridophyta of Africa, Madagascar and neighbouring islands*. South African National Biodiversity Institute, Cape Town, South Africa | 1 | 0 |
| Royal Botanic Gardens and Domain Trust (2017). PlantNET - The NSW Plant Information Network System, Available at: http://plantnet.rbgsyd.nsw.gov.au (last accessed: September 16, 2016) | 1 | 1 |
| Royal Botanic Gardens Kew (2016). Seed information database (SID). v7.1, Available at: http://data.kew.org/sid/ (last accessed: March 22, 2013) | 0 | 1 |
| Rundel, P.W., Dillon, M.O. & Palma, B. (1996). Flora and Vegetation of Pan de Azúcar National Park in the Atacama desert of Northern Chile. *Gayana Bot*, **53**, 295–315 | 1 | 1 |
| Sachet, M.-H. (1962). Flora and vegetation of Clipperton Island. *Proceedings of the California Academy of Sciences*, **31**, 249–307 | 1 | 0 |
| Sáez, L. & Rosselló, J.A. (2001). *Llibre vermell de la flora vascular de les Illes Balears*. Direcció General de Biodiversitat, Conselleria de Medi Ambient, Govern de les Illes Balears, Palma de Mallorca, Spain | 1 | 0 |
| SANBI (2014). Plants of Southern Africa. An online checklist, Available at: http://posa.sanbi.org (last accessed: March 09, 2015) | 1 | 1 |
| Sandbakk, B.E., Alsos, I.G., Arnesen, G. & Elven, R. (1996). The flora of Svalbard, Available at: http://svalbardflora.no/ (last accessed: March 16, 2011) | 1 | 0 |
| Schaefer, H., Hardy, O.J., Silva, L., Barraclough, T.G. & Savolainen, V. (2011). Testing Darwin's naturalization hypothesis in the Azores. *Ecology Letters*, **14**, 389–396 | 0 | 1 |
| Schönfelder, P. & Schönfelder, I. (1997). *Die Kosmos-Kanarenflora. Über 850 Arten der Kanarenflora und 48 tropische Ziergehölze*. Franckh-Kosmos, Stuttgart | 0 | 1 |
| Searle, J. & Madden, S. (2006). *Flora assessment of South Stradbroke Island*. Gold Coast City Council, Gold Coast City, Australia | 1 | 0 |
| Selvi, F. (2010). A critical checklist of the vascular flora of Tuscan Maremma (Grosseto province, Italy). *Fl. Medit*, **20**, 47–139 | 1 | 1 |
| Senterre, B., Chew, M.Y. & Chung, R.C.K. (2015). Flora and vegetation of Pulau Babi Tengah, Johor, Peninsular Malaysia. *Check list*, **11**, 1714 | 0 | 1 |
| Shaw, J.D., Spear, D., Greve, M. & Chown, S.L. (2010). Taxonomic homogenization and differentiation across Southern Ocean Islands differ among insects and vascular plants. *Journal of Biogeography*, **37**, 217–228 | 1 | 0 |
| Short, P.S., Albrecht, D.E., Cowie, I.D., Lewis, D.L. & Stuckey, B.M. (2011). *Checklist of the vascular plants of the Northern Territory*. Department of Natural Resources, Environment, The Arts and Sport, Darwin | 1 | 0 |
| Silaeva, T.B., Chugunov, G.G., Kiryukhin, I.V., Ageeva, A.M., Vargot, E.V., Grishutkina, G.A. & Khapugin, A.A. *Flora of the national park "Smolny". Mosses and vascular plants: annotated list of species [In Russian]*. Commission of RAS for the Conservation of Biological Diversity [Комиссия РАН по сохранению биологического разнообразия] | 1 | 0 |
| Singh, A. (2015). Observations on the vascular flora of Banaras Hindu University Main Campus, India. *International Journal of Modern Biology and Medicine*, **6**, 48–87 | 1 | 1 |
| Slik, F.J.W., Arroyo-Rodríguez, V., Aiba, S.-I., et al. (2015). An estimate of the number of tropical tree species. *Proceedings of the National Academy of Sciences of the United States of America*, **112**, 7472–7477 | 0 | 1 |
| SLUFG (2015). *Rote Liste und Artenliste Sachsen. Farn- und Samenpflanzen*. Sächsisches Landesamt für Umwelt, Landwirtschaft und Geologie, Dresden | 1 | 0 |
| Smith, A.C. (1979-1996). *Flora Vitiensis nova. a new Flora of Fiji (spermatophytes only)*. Pacific Tropical Botanical Garden (Lawaii, Hawaii) | 1 | 1 |
| Stace, C.A., Ellis, R.G., Kent, D.H. & McCosh, D.J. (2003). *Vice-county Census Catalogue of the vascular plants of Great Britain, the Isle of Man and the Channel Islands*. Botanical Society of the British Isles, London, UK | 1 | 0 |
| Stalmans, M. (2006). *Tinley's plant species list for the Greater Gorongosa ecosystem, Moçambique*. Unpublished report by International Conservation Services to the Carr Foundation and the Ministry of Tourism | 1 | 1 |
| Stalter, R. & Lamont, E.E. (2006). The historical and extant flora of Sable Island, Nova Scotia, Canada. *Journal of the Torrey Botanical Society*, **133**, 362–374 | 1 | 0 |
| Stoddart, D.R. & Fosberg, F.R. (1994). Flora of the Phoenix Islands, central Pacific. *Atoll Research Bulletin*, **393**, 1–60 | 1 | 0 |
| Strahm, W.A. *The conservation and restoration of the flora of Mauritius and Rodrigues* | 1 | 0 |
| Sykes, W.R. (1970). Contributions to the flora of Niue. *Bulletin. Department of Scientific and Industrial Research, New Zealand*, **200**, 321 | 1 | 0 |
| Sykes, W.R., West, C.J., Beever, J.E. & Fife, A.J. (2000). *Kermadec Islands flora-special edition. a compilation of modern material about the flora of the Kermadec Islands*. Manaaki Whenua Press, Landcare Research | 1 | 0 |
| Takahashi, H., Barkalov, V.Y., Gage, S. & Zhuravlev, Y.N. (1997). A preliminary study of the flora of Chirpoi, Kuril Islands. *Acta Phytotaxonomica et Geobotanica*, **48**, 31–42 | 1 | 0 |
| Takahashi, H., Barkalov, V.Y., Gage, S., Joneson, S., Ilushko, M. & Zhuravlev, Y.N. (2002). A floristic study of the vascular plants of Raikoke, Kuril Islands. *Acta Phytotaxonomica et Geobotanica*, **53**, 17–33 | 1 | 0 |
| Takahashi, H., Barkalov, V.Y., Gage, S., Semsrott, B., Ilushko, M. & Zhuravlev, Y.N. (1999). A preliminary checklist of the vascular plants of Chirinkotan, Kuril Islands. *Journal of Phytogeography and Taxonomy*, **47**, 131–137 | 1 | 0 |
| Takahashi, H., Barkalov, V.Y., Gage, S., Semsrott, B., Ilushko, M. & Zhuravlev, Y.N. (2006). A floristic study of the vascular plants of Kharimkotan, Kuril Islands. *Bulletin of the Hokkaido University Museum*, **3**, 41–66 | 1 | 0 |
| Tamis, W.L.M., van der Meijden, R., Runhaar, J., Bekker, R.M., Ozinga, W.A., Odé, B. & Hoste, I. (2004). Standard List of the Flora of the Netherlands 2003. *Gorteria*, **30**, 101–195 | 1 | 1 |
| Tatewaki, M. (1957). Geobotanical studies on the Kurile Islands. *Acta Horti Gotoburgensis*, **21**, 43–123 | 1 | 0 |
| Taylor, R. (2006). *Straight through from London. the Antipodes and Bounty Islands, New Zealand*. Heritage Expeditions New Zealand, Christchurch, New Zealand | 1 | 0 |
| Tela Botanica (2015). Base de données des Trachéophytes de France métropolitaine (bdtfx), Available at: http://www.tela-botanica.org/bdtfx | 1 | 0 |
| Ter Steege, H., Vaessen, R.W., Cardenas-Lopez, D., Sabatier, D., Antonelli, A., Oliveira, S.M. de, Pitman, N.C.A., Jorgensen, P.M. & Salomao, R.P. (2016). The discovery of the Amazonian tree flora with an updated checklist of all known tree taxa. *Scientific reports*, **6**, 29549 | 0 | 1 |
| Thaman, R.R., Fosberg, F.R., Manner, H.I. & Hassall, D.C. (1994). The flora of Nauru. *Atoll Research Bulletin*, **392**, 1–233 | 1 | 0 |
| Thomas, J. (2011). Plant diversity of Saudi Arabia. Flora checklist, Available at: http://plantdiversityofsaudi-arabia.info/biodiversity-saudi-arabia/flora/Checklist/Checklist.htm (last accessed: January 14, 2016) | 1 | 0 |
| Tonkov, S., Pavlova, D., Atanassova, J., Nedelcheva, A. & Marinova, E. (2004). *Floristic catalogue of the nature reserve Rilomanastirska Gora (Central Rila Mountains). I. The locality Kirilova*. University of Sofia | 0 | 1 |
| Tropicos (2015). Catálogo de las Plantas Vasculares de Bolivia. *Tropicos*. Missouri Botanical Garden, St. Louis, Available at: http://www.tropicos.org/Project/BC | 1 | 1 |
| Tropicos (2015). Catalogue of the Vascular Plants of Ecuador, Available at: http://www.tropicos.org/Project/CE (last accessed: October 22, 2015) | 1 | 1 |
| Tropicos (2015). Catalogue of the Vascular Plants of the Department of Antioquia (Colombia). *Tropicos*. Missouri Botanical Garden, St. Louis, Available at: http://www.tropicos.org/Project/CV | 0 | 1 |
| Tropicos (2015). Flora de Nicaragua. *Tropicos*. Missouri Botanical Garden, St. Louis, Available at: http://www.tropicos.org/Project/FN | 1 | 0 |
| Tropicos (2015). Panama Checklist. *Tropicos*. Missouri Botanical Garden, St. Louis, Available at: http://www.tropicos.org/Project/PAC | 0 | 1 |
| Tropicos (2015). Paraguay Checklist. *Tropicos*. Missouri Botanical Garden, St. Louis, Available at: http://www.tropicos.org/Project/Paraguay | 0 | 1 |
| Tropicos (2015). Peru Checklist. *Tropicos*. Missouri Botanical Garden, St. Louis, Available at: http://www.tropicos.org/Project/PEC | 0 | 1 |
| Trusty, J.L., Kesler, H.C. & Delgado, G.H. (2006). Vascular flora of Isla del Coco, Costa Rica. *Proceedings of the California Academy of Sciences*, **57**, 247–355 | 1 | 0 |
| Turner, I.M. (1995). *A catalogue of the vascular plants of Malaya*. Gardens' Bulletin, Singapore | 1 | 1 |
| Tutul, E., Uddin, M.Z., Rahman, M.O. & Hassan, M.A. (2009). Angiospermic flora of Runtia sal forest, Bangladesh. I. Liliopsida (Monocots). *Bangladesh Journal of Plant Taxonomy*, **16**, 83–90 | 0 | 1 |
| UIB (2007). Herbario virtual del Mediterráneo Occidental, Available at: http://herbarivirtual.uib.es/cas-med/ (last accessed: August 07, 2012) | 1 | 0 |
| University of Greifswald (2010). FloraGREIF - Virtual Flora of Mongolia, Available at: http://greif.uni-greifswald.de/floragreif/ (last accessed: February 02, 2016) | 0 | 1 |
| University of Kent (2012). Cook Islands Biodiversity and Ethnobiology Database, Available at: http://cookislands.pacificbiodiversity.net/cibed/dbs/search.html (last accessed: April 12, 2012) | 1 | 0 |
| USDA & NRCS (2015). The PLANTS Database. National Plant Data Team (ed.), Available at: http://plants.usda.gov (last accessed: April 24, 2015) | 1 | 1 |
| van Vreeswyk, A.M.E., Payne, A.L., Leighton, K.A. & Hennig, P. (2004). *An inventory and condition survey of the Pilbara region, Western Australia*. Department of Agriculture | 0 | 1 |
| Vanderplank, S.E. (2010). *The Vascular Flora of Greater San Quintín, Baja California, Mexico*. CGU Theses & Dissertations | 1 | 1 |
| Velarde, E., Wilder, B.T., Felcer, R.S. & Ezcurra, E. (2014). Floristic diversity and dynamics of Isla Rasa, Gulf of California - A globally important seabird island. *Botanical Sciences*, **92**, 89–101 | 0 | 1 |
| Velarde, E.P., Guzmán, R.C. & Koch, S.D. (2008). Plantas vasculares y vegetación de la parte alta del Arroyo Agua Fría, municipio de Minatitlán, Colima, México. *Acta Botanica Mexicana*, **84**, 25–72 | 0 | 1 |
| Velayos Rodríguez, M. (2016). Flora de Guinea Equatorial, Available at: http://www.floradeguinea.com/ (last accessed: February 14, 2016) | 1 | 0 |
| Velayos, M., Barberá, P., Cabezas, F.J., La Estrella, M.d., Fero, M. & Aedo, C. (2014). Checklist of the Vascular Plants of Annobón (Equatorial Guinea). *Phytotaxa*, **171**, 1–78 | 1 | 0 |
| VicFlora (2016). Flora of Victoria, Available at: http://vicflora.rbg.vic.gov.au (last accessed: November 30, 2016) | 1 | 0 |
| Viciani, D., Gonnelli, V., Sirotti, M. & Agostini, N. (2010). An annotated check-list of the vascular flora of the “Parco Nazionale delle Foreste Casentinesi, Monte Falterona e Campigna”(Northern Apennines Central Italy). *Webbia*, **65**, 3–131 | 1 | 1 |
| Vogt, C. (2011). Composición de la Flora Vascular del Chaco Boreal, Paraguay. I. Pteridophyta y Monocotiledoneae. *Steviana*, **3**, 13–47 | 1 | 1 |
| Wace, N.M. & Dickson, J.H. (1965). The terrestrial botany of the Tristan da Cunha Islands. *Philosophical Transactions of the Royal Society of London B Biological Sciences*, **249**, 273–360 | 1 | 0 |
| Wace, N.M. (1961). The vegetation of Gough Island. *Ecological Monographs*, **31**, 337–367 | 1 | 0 |
| Wagner, W.L. & Lorence, D.H. (2002). Flora of the Marquesas Islands website, Available at: http://botany.si.edu/pacificislandbiodiversity/marquesasflora/index.htm (last accessed: November 28, 2014) | 1 | 0 |
| Wagner, W.L., Herbst, D.R. & Lorence, D.H. (2005). Flora of the Hawaiian Islands website, Available at: http://botany.si.edu/pacificislandbiodiversity/hawaiianflora/ (last accessed: October 16, 2010) | 1 | 1 |
| WCSP (2014). World Checklist of Selected Plant Families, Available at: http://apps.kew.org/wcsp/home.do (last accessed: December 01, 2014) | 1 | 0 |
| Webster, G.L. & Rhode, R.M. (2001). Plant diversity of an Andean cloud forest: inventory of the vascular plants of Maquipucuna, Ecuador. *Publications in Botany*, **82**, 1–228 | 0 | 1 |
| Wellington Botanical Society (2008). *Native vascular plants of Great Barrier Island*. Wellington Botanical Society, Wellington, New Zealand | 1 | 0 |
| Wester, L. (1985). Checklist of the vascular plants of the northern Line Islands. *Atoll Research Bulletin*, **187**, 1–38 | 1 | 0 |
| Western Australian Herbarium (2017). FloraBase - the Western Australian Flora. Western Australian Herbarium (ed.), Available at: https://florabase.dpaw.wa.gov.au/ (last accessed: September 16, 2016) | 1 | 1 |
| Whistler, W.A. (1983). Vegetation and flora of the Aleipata Islands, Western Samoa. *Pacific Science*, **37**, 227–249 | 1 | 0 |
| Whistler, W.A. (1998). *A study of the rare plants of American Samoa*. US Fish and Wildlife Service, Honolulu, Hawaii | 1 | 0 |
| Whistler, W.A. (2012). Botanical survey of the Ringgold Islands, Fiji. *Allertonia*, **11**, 1–28 | 1 | 0 |
| Whistler, W.A. *Botanical survey of Diego Garcia, Chagos Archipelago, British Indian Ocean Territory*. Appendix E1 | 1 | 0 |
| Woodroffe, C.D. (1986). Vascular plant species-area relationships on Nui Atoll, Tuvalu, Central Pacific: a reassessment of the small island effect. *Australian Journal of Ecology*, **11**, 21–31 | 1 | 1 |
| Yineger, H., Kelbessa, E., Bekele, T. & Lulekal, E. (2008). Floristic composition and structure of the dry afro-montane forest at Bale Mountains National Park, Ethiopia. *SINET: Ethiopian Journal of Science*, **31**, 103–120 | 0 | 1 |
| ZDSF & SKEW (2014). Info Flora. Artenliste Schweiz 5x5 km, Available at: https://www.infoflora.ch/de/daten-beziehen/artenliste-5x5-km.html (last accessed: February 13, 2015) | 1 | 0 |
| Zhang, S.T., Zhen Du, G. & Chen, J.K. (2004). Seed size in relation to phylogeny, growth form and longevity in a subalpine meadow on the east of the Tibetan plateau. *Folia Geobotanica*, **39**, 129–142 | 0 | 1 |
| Zuloaga, F.O., Morrone, O. & Belgrano, M. (2014). Catálogo de las Plantas Vasculares del Cono Sur, Available at: http://www.darwin.edu.ar/Proyectos/FloraArgentina/fa.htm (last accessed: March 16, 2015) | 1 | 1 |
| Zvyagintseva, K.O. (2015). *An annotated checklist of the urban flora of Kharkiv*. Kharkiv National University, Kharkiv, Ukraine | 1 | 0 |
| Артемов, И.А. (2012). *Определитель растений Катунского биосферного заповедника*. Russian Academy of Sciences, БАРНАУЛ | 0 | 1 |
| Евстигнеев, О.И. & Федотов, Ю.П. (2007). *Флора сосудистых растений заповедника "Брянский лес"*. Гос. природ. биосфер. заповедник Брян. лес, Брянск | 1 | 0 |
| Хапугин, А.А. (2013). *Сосудистые растения Ромодановского района Республики Мордовия (конспект флоры)*, Saransk | 1 | 0 |

