## Supplementary material for "Disharmony of the world’s island floras": SI_items (from another version of this preprint: version 2, 2019-03-26)

### Supporting information

**SI Figures**

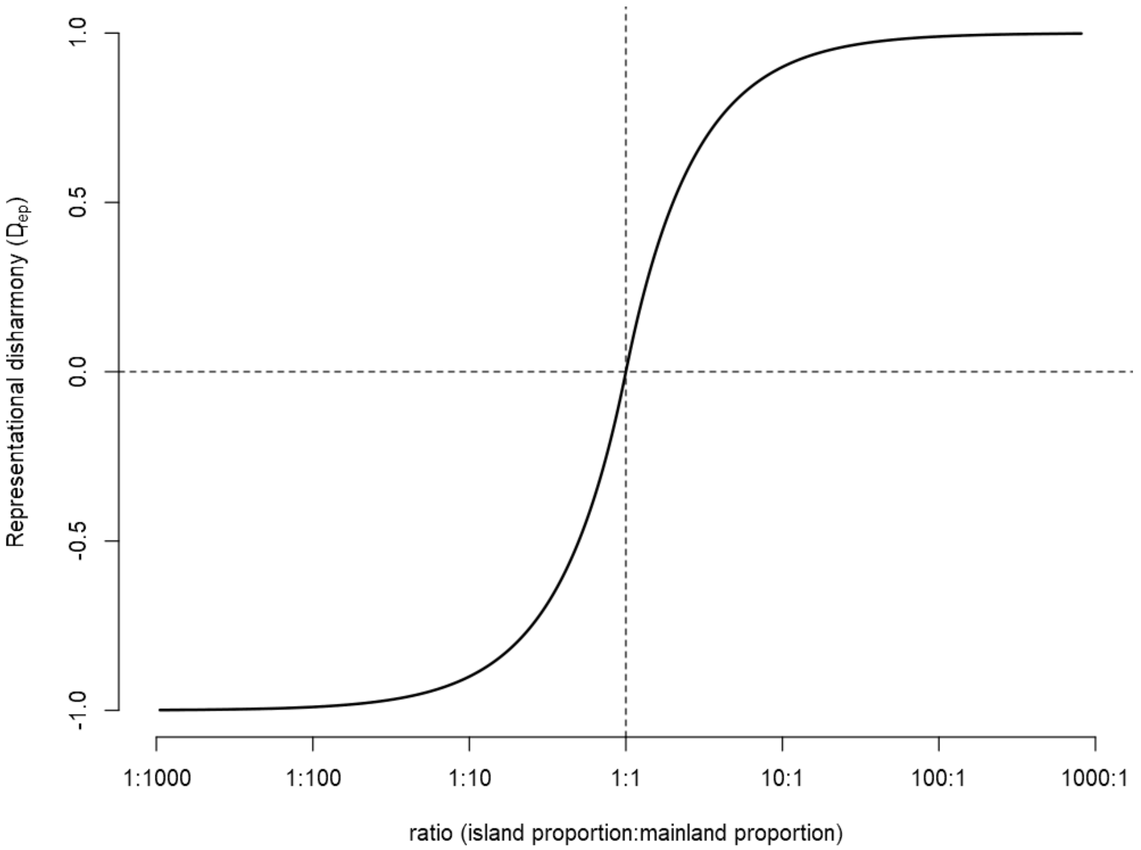


*SI Figure 1: Representational disharmony index ($D_{rep}$) as a function of the ratio between mean proportional representation in island- and mainland floras.*

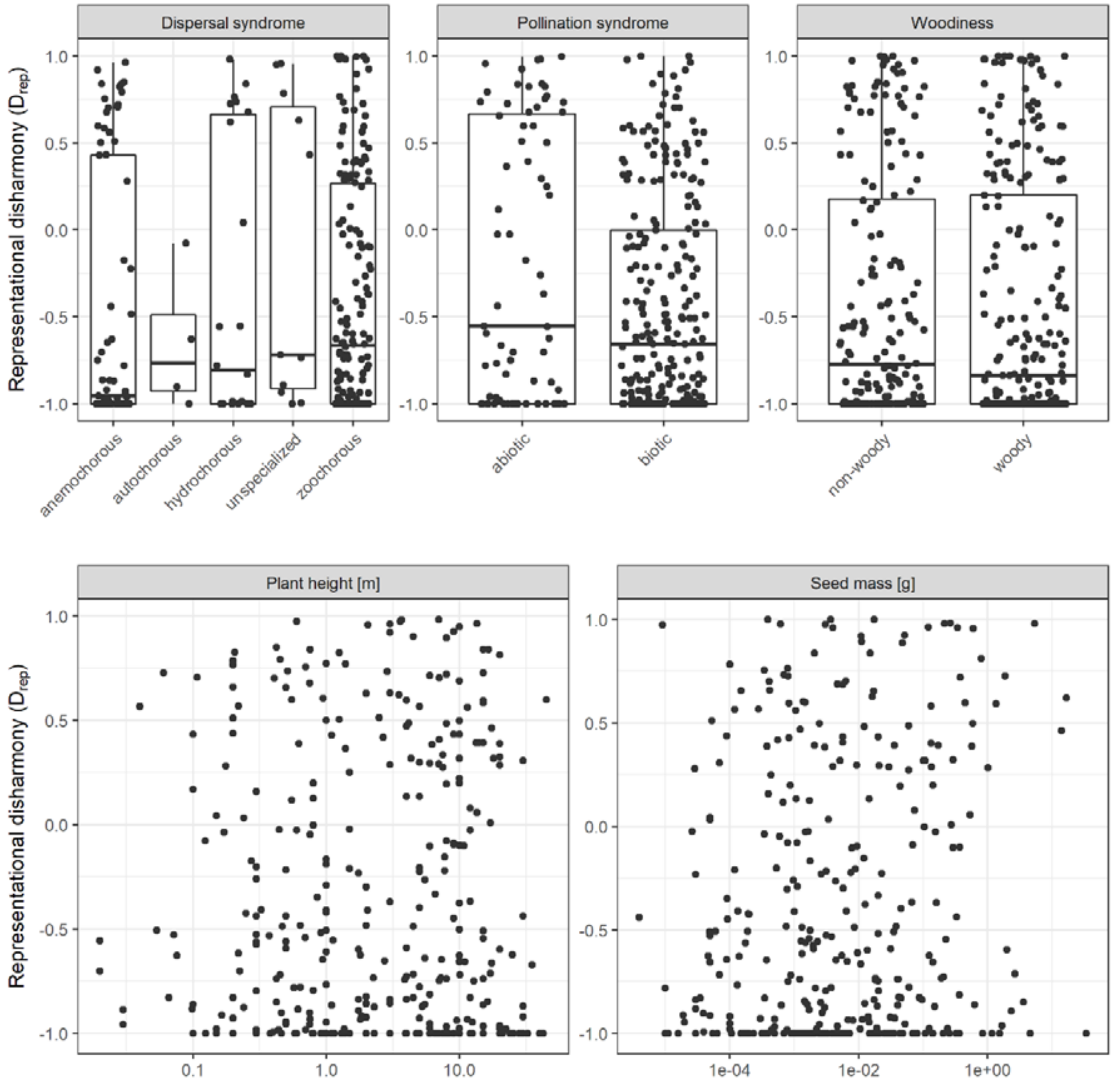


*SI Figure 2: Relationship between representational disharmony ($D_{rep}$) and family-level functional traits.*

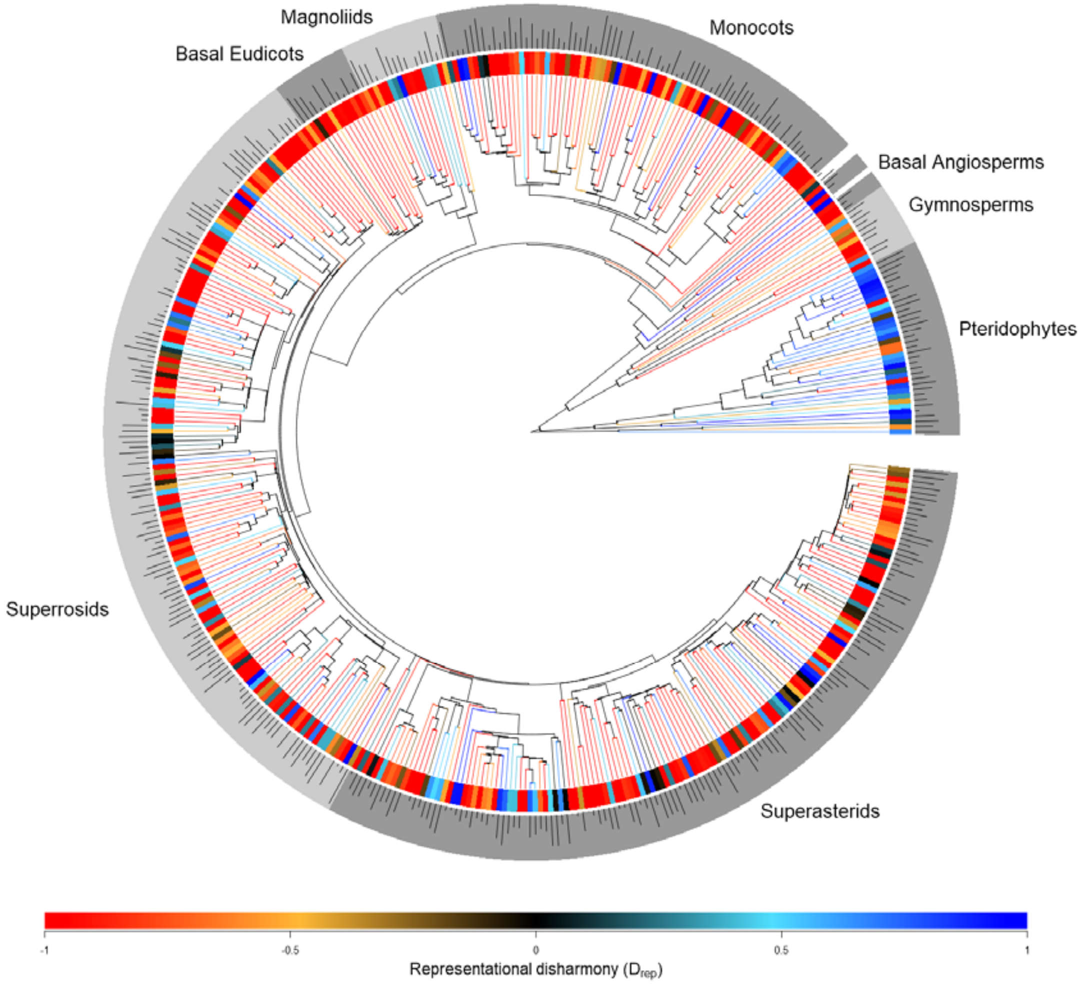


*SI Figure 3: Representational disharmony ($D_{rep}$) of 450 vascular plant families. Negative values indicate a proportional under-representation of the respective family in island floras compared to the mainland, whereas positive values indicate an over-representation. $D_{rep} = -1$, $D_{rep} = 0$ and $D_{rep} = 1$ respectively denote a restriction to the mainland, even representation in island and mainland floras, and a restriction to islands.*

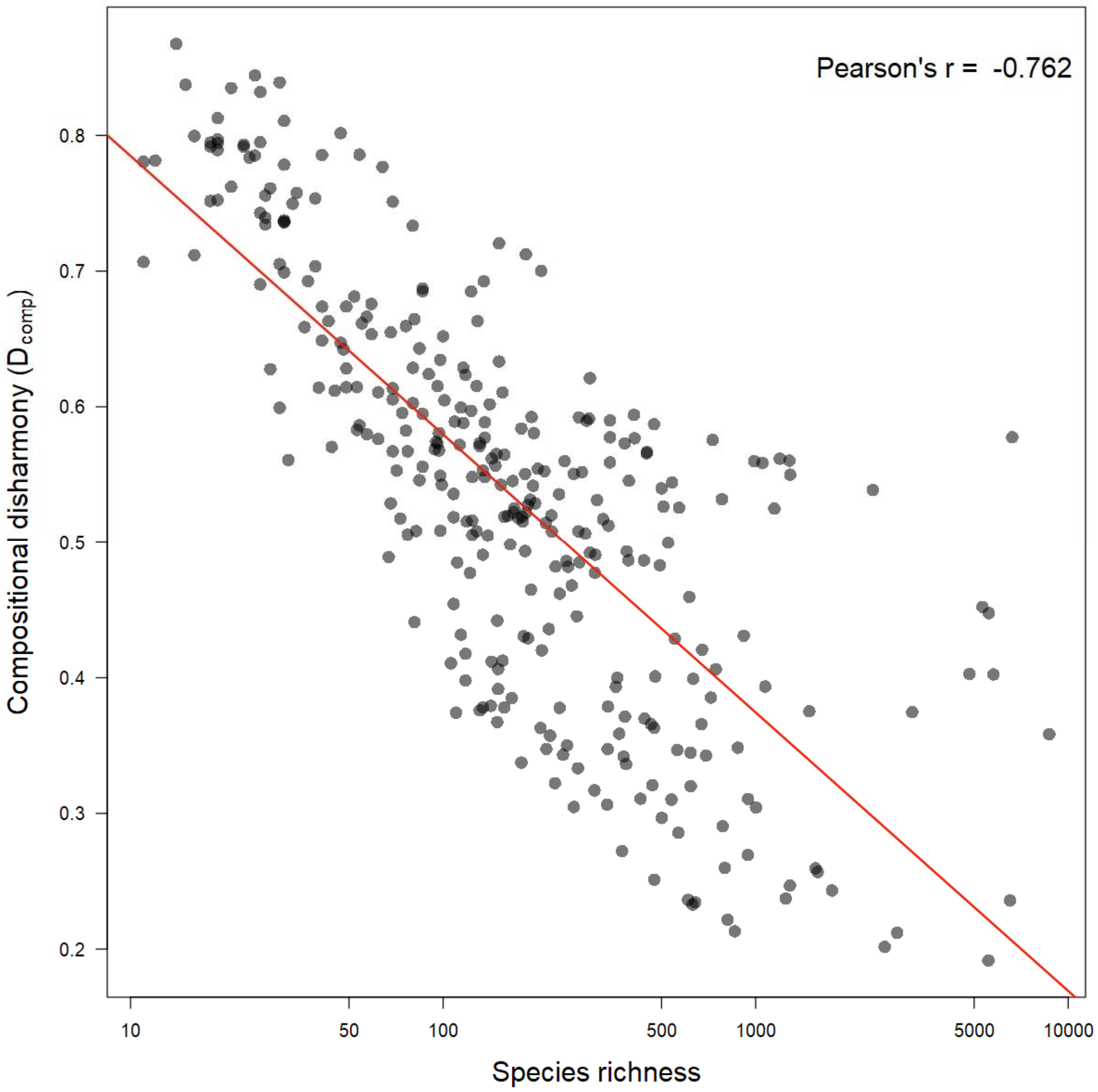


*SI Figure 4: Correlation between compositional disharmony ($D_{comp}$) and $log_{10}$(species richness).*

**SI Text 1: Truncation of mainland weights vector $W$**

The presented measure of compositional disharmony is calculated as the mean Bray-Curtis dissimilarity of family proportions of a given island $i$ relative to all mainland floras ($B_i$), weighted by the mainland floras' estimated importance as source regions for the respective island ($W_i$):

$$D_{comp}(i) = \frac{\sum B_i * W_i}{\sum W_i}$$

We obtained $W_i$ from predictions of species turnover between island $i$ and all mainland units. The model we used to predict species turnover (Generalized dissimilarity modelling, Ferrier 2002, Ferrier et al. 2007) converges against, but never actually reaches 1 (complete species turnover) at large geographical and/or environmental distances. This is generally a realistic and useful property, because even remote or environmentally dissimilar regions might share a small fraction of their species. However, in our particular use case, the combined weight of many mainland regions with relatively low values of $W_i$ could still introduce a substantial bias into $D_{comp}$. This problem was especially pronounced for islands having only very few important mainland source regions, e.g. due to the lack of floristic data for nearby continental regions in our dataset.

To avoid a dependency of $D_{comp}$ on the amount and geographical distribution of available mainland floras, we set $W_i = 0$ for mainland units falling below a certain island-specific value. To determine this threshold, we performed the following steps:

1. Scale the source region weights vector $W_i$ for a given island $i$ to a range of 0 to 1.
2. Sort $W_i$ in decreasing order, i.e. rank all mainland units according to their importance as source region for island i (SI Figure 5, black line).
3. Fit a curve to the ordered values in $W_i$ to obtain a smooth and continuous approximation of the weights distribution (SI Figure 5, blue line). We used the `smooth.spline`-function in R with a smoothing parameter of `spar = 0.75`.
4. Take the derivative $W_i'$ to quantify the rate of change in $W_i$.
5. Define the threshold for island i as that value in $W_i$, after which $W_i'$ permanently falls below $0.001$, i.e. no more substantial changes in the importance of mainland source regions are predicted (SI Figure 5, red lines). The value of 0.001 was chosen after carefully inspecting curves of $W_i$ and $W_i'$ for a representative set of island floras.

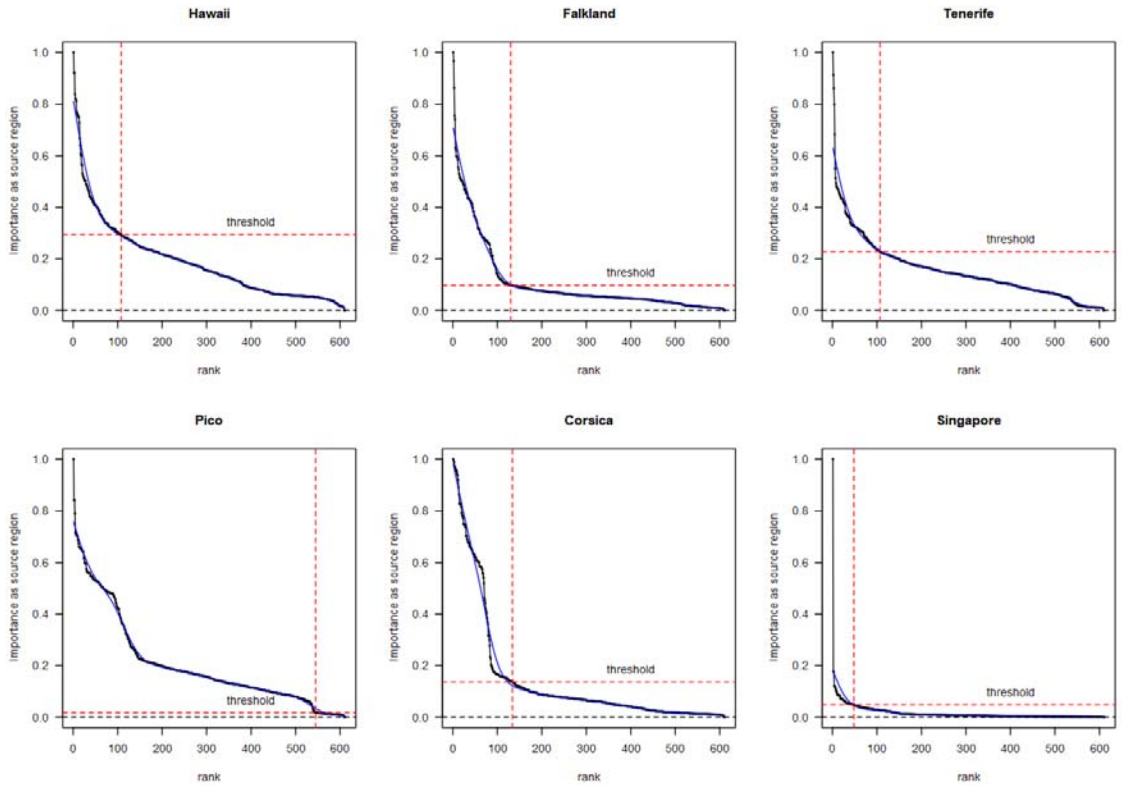


SI Figure 5: Determination of threshold value for weights vector W, exemplarily for six islands.
